# Metabolic remodeling by TIGAR overexpression is a therapeutic target in esophageal squamous-cell carcinoma

**DOI:** 10.1101/789271

**Authors:** Jiahui Chu, Xiangjie Niu, Jiang Chang, Mingming Shao, Linna Peng, Yiyi Xi, Ai Lin, Chengcheng Wang, Qionghua Cui, Yingying Luo, Wenyi Fan, Yamei Chen, Yanxia Sun, Wenjia Guo, Wen Tan, Dongxin Lin, Chen Wu

## Abstract

Whole-genome sequencing has identified many amplified genes in esophageal squamous-cell carcinoma (ESCC); however, their roles and the clinical relevance have yet elucidated. Here we show TP53-induced glycolysis and apoptosis regulator (TIGAR) is a major player in ESCC progression and chemoresistance. TIGAR reprograms glucose metabolism from glycolysis to the glutamine pathway through AMP-activated kinase, and its overexpression is correlated with poor disease outcomes. *Tigar* knockout mice have reduced ESCC growth and tumor burden. Treatment of TIGAR-overexpressed ESCC cell xenografts and patient-derived tumor xenografts in mice with combination of glutaminase inhibitor and chemotherapeutic agents achieves significant more efficacy than chemotherapy alone. These findings shed light on an important role of TIGAR in ESCC and provide evidence for targeted treatment of TIGAR-overexpressed ESCC.

**Significance:** Effective and target therapies are required for ESCC, one of the most common types of digestive OR cancer. Little has been known about the biology of ESCC progression or potential molecular targets OR for treatment. Whole-genome sequencing and RNA sequencing studies in ESCC have identified OR many recurrent copy number gain genes; however, the roles and druggable relevance of these OR genes remains poorly understood. Herein we demonstrate that TIGAR overexpression leads to OR metabolic remodeling, promoting ESCC progression and resistance to chemotherapeutic agents. OR Inhibiting the glutamine pathway significantly represses TIGAR-overexpressing ESCC growth and OR enhances tumor cell sensitization to cytotoxic agents. These findings might provide the rationale OR for clinical trials testing glutamine pathway inhibitors in combination with chemotherapy in OR TIGAR-expressing ESCC.

## INTRODUCTION

Esophageal squamous cell carcinoma (ESCC) is one of the leading causes of cancer-related mortality in China and some other parts of the world (Chen et al., 2016; Pennathur et al., 2013; Siegel et al., 2016). Currently, surgery is the major approach to treat ESCC but only a portion of patients with ESCC has the chance to be treated with esophagectomy (Ohashi et al., 2015; Wouters et al., 2012). For locally advanced ESCC, chemotherapy and radiotherapy are commonly used but the survival rates of patients receiving the remedies are dismal (Bedenne et al., 2007; Crosby et al., 2013; Stahl et al., 2005). ESCC lacks effective targeted therapies mainly because we know little about the biology of its progression or potential molecular targets for its treatment. Therefore, better understanding the molecular mechanism of ESCC progression and seeking more effective drug targets are extremely needed.

In recent years, high-throughput genome-wide screening for amplified and overexpressed genes has accelerated the discovery of potential molecular targets for drug development (Guichard et al., 2012; Santarius et al., 2010). In our previous whole-genome sequencing (WGS) study on 94 ESCC samples, we identified 23 focal recurrent copy number gain regions containing 1,591 genes and the matched mRNA expression data showed 149 copy-number gain genes were overexpressed in tumor compared with adjacent normal samples (Chang et al., 2017). These results are consistent with the TCGA ESCC data, indicating that ESCC is a type of cancer dominated by CNVs (Beroukhim et al., 2010). In the present study, we further investigated the relevance of previously identified 149 CNV genes in the proliferation and progression of ESCC to search for potential treatment targets. To distinguish the genomic alterations that drive cancer growth and progression from numerous passenger alterations accumulating during tumorigenesis, we employed RNA interfering-based high-content screening followed by a series of functional analyses to evaluate the roles of candidate genes in the malignant phenotypes of ESCC cells. In this effort, we identified TP53-induced glycolysis and apoptosis regulator (TIGAR) as an important player.

Previous studies have shown that TIGAR may function as fructose-2,6-bisphosphatase that degrades intracellular fructose-2,6-bisphosphate, which dampens glycolysis by reducing the phosphofrutokanase-1 activity (Bensaad et al., 2006; Okar and Lange, 1999; Rider et al., 2004). It has been proposed that glycolysis inhibition by TIGAR may divert the glycolytic flux to alternative metabolic pathways including the pentose phosphate pathway that generates more reductants such as NADPH and GSH to protect cancer cells from reactive oxygen species, and the hexosamine pathway that promotes nucleotide synthesis for cancer cell proliferation (Bensaad et al., 2006; Yang et al., 2017). The overexpression of TIGAR has been associated with some types of human cancer (Lee et al., 2014). However, little has been known about TIGAR and its molecular function and clinic relevance in ESCC.

Here, we report the oncogenic role of TIGAR in ESCC progression and resistance to chemotherapy. We found that TIGAR is overexpressed in most of human ESCC samples and presents as an early molecular event in 4-nitroquinoline N-oxide (4-NQO)-induced mice ESCC. Overexpressed TIGAR remodels energy metabolism from glycolysis to the glutamine pathway through the activation of AMP-activated protein kinase (AMPK), resulting in glutamine addition in the progression of ESCC. We also found that treatment of TIGAR-overexpressing ESCC cell xenografts and patient-derived xenografts (PDXs) in mice with combination of glutaminase inhibitor and cytotoxic chemotherapeutic agents has significantly more effective than treatment with chemotherapy alone.

## Results

### TIGAR is an important player in ESCC proliferation and progression

Using RNA interfering-based high content screening in two ESCC cell lines KYSE450 and KYSE510 (Figure 1A), we found that knockdown of 18 of the 149 select candidate genes (see Methods for detail; Figure S1) had significant inhibitory effects on cancer cell proliferation (P<0.05) (Figure 1B). Among the genes, knockdown of TIGAR expression most efficiently suppressed the malignant phenotypes in ESCC cells, with the cell-proliferation inhibition rates of 44.2% (P=0.002) in KYSE450 and 40.3% (P=0.002) in KYSE510 compared with controls (Figure 1B). We then established ESCC cell lines (KYSE150 and KYSE30) with *TIGAR* knockout or stable overexpression to verify this gene effects on cancer cell growth in vitro and in vivo in mice. We found that *TIGAR* knockout in KYSE150 and KYSE30 cells significantly repressed these cell growth, whereas the ectopic *TIGAR* overexpression significantly promoted proliferation (Figures 1C–1F and Figure S2A). Such effects also existed in vivo when these cells were subcutaneously transplanted into the armpit of nude mice: the xenograft tumors derived from TIGAR-overexpressing ESCC cells had significantly higher growth rates than controls transfected with blank vector (Figure 1G).

**Figure 1.**
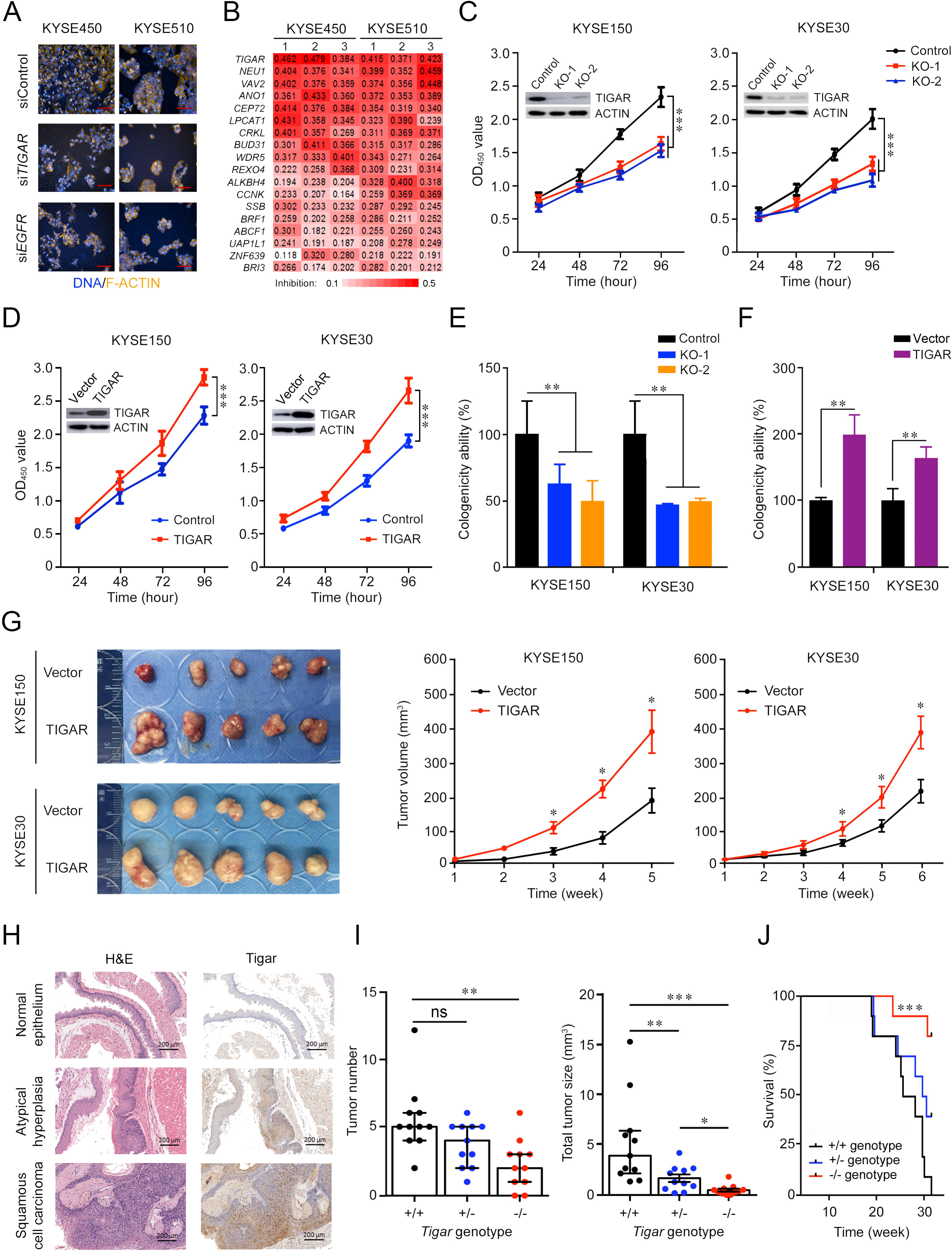
Identification of TIGAR as a player in ESCC proliferation and progression. (A) High-content phenotyping of ESCC cells transfected with siRNA targeting *TIGAR*. Cells transfected with scramble siRNA (siContol) or siRNA targeting *EGFR* served as negative or positive controls, respectively. (B) Heat map displaying inhibitory effects of knockdown of 18 genes in two ESCC cell lines with 3 replications. (C–F) TIGAR knockout (KO) significantly suppressed but TIGAR overexpression significantly promoted ESCC cell proliferation (C and D) and colony formation (E and F). Error bars represent SEM obtained from three independent experiments. **, *P*<0.01 and ***, *P*<0.001 from Student’s *t*-test. (G) TIGAR overexpression promoted tumor xenograft growth in vivo in nude mice. *Left* panel shows tumors in the end of the experiment and *right* plots show tumor growth curves measured every 7 days after injection of ESCC cells with or without TIGAR overexpression. Tumor volume = 0.5 × length × width^2^. Error bars represent SEM. *, *P*<0.05 from Mann–Whitney test. (H) Hematoxylin and eosin (H&E) staining (*left*) and immunohistochemistry analysis of Tigar (*right*) in mice primary ESCC induced by 4-NQO (see Methods for details) at the different stages, showing that Tigar overexpression is an early event in ESCC development and plays an important role in ESCC progression. (I) Effects of *Tigar* knockout on ESCC formation and tumor burden induced by 4-NQO in mice. *Left* panel shows tumor number and *right* panel shows total tumor size in mice with *Tigar*^+/+^, *Tigar*^+/-^ or *Tigar*^-/-^ genotype. Error bars represent SEM. *, P<0.05; **, P<0.01 and ***, P<0.001 of Mann-Whitney test. (J) Kaplan-Meier estimate of survival among mice with *Tigar*^+/+^, *Tigar*^+/-^ or *Tigar*^-/-^ genotype receiving 4-NQO. ***, P<0.001 of log-rank test.

We then performed ESCC tumorigenesis induced by chemical carcinogen 4-NQO in mice with or without *Tigar* knockout (Figures S2B and S2C). Esophageal atypical hyperplasia lesion and squamous cell carcinoma could be identified at week 20 and 24, respectively. IHC staining of *Tigar* protein in the esophageal tissue samples from *Tigar*^+/+^ mice showed substantially high *Tigar* expression in atypical hyperplasia lesion and ESCC than in normal esophageal epithelium (Figure 1H), suggesting that *Tigar* overexpression might play an important role in the development and progression of ESCC. At the experiment end of 30 weeks, all mice with the *Tigar*^+/+^ or *Tigar*^+/-^ genotype (11/11) developed ESCC, but only 82% (9/11) of *Tigar*^-/-^ mice had ESCC at the same time (Figure S2D). Furthermore, *Tigar*^-/-^ mice had a significantly smaller median tumor number and median total tumor size per animal (2 and 0.39, respectively) than *Tigar*^+/+^ (5 and 3.90, respectively) or *Tigar*^+/-^ (4 and 1.30, respectively) mice (Figure 1I). Accordingly, we observed a significantly longer survival time in *Tigar*^-/-^ mice than in *Tigar*^+/+^ or *Tigar*^+/-^ mice treated with the carcinogen (Figure 1J).

### TIGAR overexpression is correlated with advanced stages and metastasis of human ESCC

We next examined the TIGAR protein expression by IHC staining in tissue arrays derived from surgically removed 225 ESCC samples and found that 177 (78.6%) of the samples had significantly higher TIGAR expression (as measured by the IHC score) than the corresponding adjacent normal tissues (*P*<1.93e-33) (Figure 2A). This result was verified in randomly selected 12 fresh ESCC and paired normal tissue samples by Western blot analysis (Figure 2B). TIGAR IHC score was significantly higher (*P*<9.30e-6) in tumor at advanced stages (III and IV) than in tumors at early stages (I and II) (Fig. 2C) and in tumors with lymph node metastasis than in tumors without lymph node metastasis (*P*<1.12e-5) (Figure 2D). Patients with high TIGAR level in ESCC survived significantly shorter time than patients with low TIGAR level in ESCC (log-rank *P*=5.25e-12; hazard ratio=4.11; 95% confidence interval=2.75–6.16) (Figure 2E). We obtained consistent results using in vitro assays: TIGAR knockout significantly suppressed while TIGAR overexpression significantly promoted the migration and invasion abilities of ESCC cells (Figures 2F and 2G; Figures S3A and S3B). These results clearly indicate that TIGAR overexpression in ESCC promotes tumor progression and is related to the disease outcomes.

**Figure 2.**
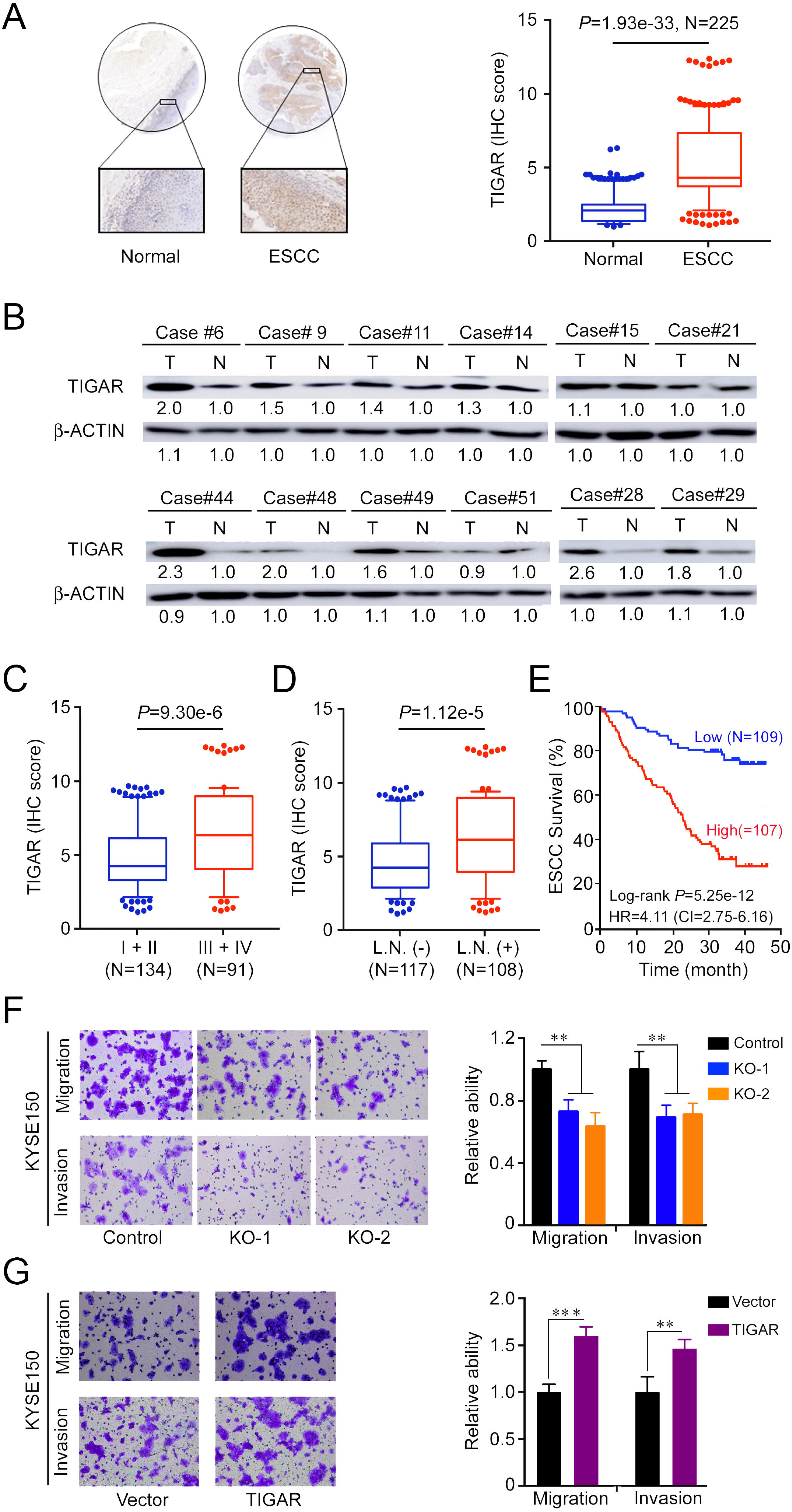
TIGAR is overexpressed in human ESCC and correlated with tumor stages and metastasis. (A) Immunohistochemistry (IHC) analysis of TIGAR expression levels in paired human ESCC and adjacent normal tissues. *Left* panel shows representative IHC pictures of tissue arrays (scale bar, 40 μm) and *right* panel shows IHC scores indicating that TIGAR expression levels were significantly higher in ESCC tumors compared with normal tissues. *P* values are from Mann-Whitney test. (B) Western blot analysis of TIGAR protein levels in randomly selected 12 paired ESCC and adjacent normal tissues, showing that most of ESCC expressed higher TIGAR than their normal tissues. (C) TIGAR protein levels were significantly higher in advanced ESCC (stages III and IV) than in early ESCC (stages I and II). *P* values are from Mann-Whitney test. (D) TIGAR protein levels were significantly higher in ESCC with lymph node metastasis than in ESCC without lymph node metastasis. *P* values are from Mann-Whitney test. (E) Kaplan-Meier estimates of survival time in 216 patients with ESCC by different TIGAR levels in tumor tissues expressed as IHC scores. (F and G) TIGAR knockout (KO) significantly suppressed the migration and invasion abilities of KYSE150 cells (F). TIGAR overexpression significantly promoted the migration and invasion abilities of KYSE150 cells (G). *Left* panels are representative pictures of ESCC cells in transwell assays and *right* panels represent data (mean ± SEM) from 3 independent experiments and each had duplicate. **, *P*<0.01 and ***, *P*<0.001 of t-test.

### TIGAR inhibits glycolysis but activates glutamine pathway to promote ESCC

We next sought to understand why TIGAR overexpression promotes ESCC proliferation and progression. Previous report has shown that TIGAR acts as a fructose-2,6-bisphosphatase that inhibits glycolysis and promoting the pentose phosphate pathway (Bensaad et al., 2006; Okar and Lange, 1999; Rider et al., 2004). We therefore examined the lactate levels and NADPH to NADP ratio in ESCC cells and found that TIGAR overexpression greatly reduced lactate but increased NADPH production in both KYSE150 and KYSE30 cells (Figures 3A and 3B). Since glycolysis is the major and necessary energy metabolism maintaining the proliferation of cancer cells, the inhibition of glycolysis by TIGAR overexpression in ESCC cells hints the existence of a compensatory energy-providing pathway. This notion is confirmed by comparing the ATP levels in TIGAR-overexpressing cells with control cells: the ATP levels in TIGAR-overexpressing cells markedly increased rather than decreased (Figure 3C). In pursuit of another metabolism pathway that sustains ESCC cell progression, we focused on the glutamine pathway since glutamine addiction has been shown to play a crucial role in many rapidly proliferating cells including cancer cells (Wise and Thompson, 2010; Yang et al., 2017). We found that the RNA and protein levels of glutamine pathway genes *SLC1A5, GLS* and *GLUD1* significantly increased in ESCC cells with TIGAR overexpression but declined in cells with TIGAR knockout as compared with control cells (Figures 3D and 3E; Figures S4A-S4D). Interestingly, TIGAR overexpression resulted in substantial AMPK activation (Thr172 phosphorylation) and GLS upregulation in a time-dependent manner (Figure 3F). AMPK expression knockdown by siRNA or inhibition of phosphorylated AMPK (p-AMPK) activity by specific inhibitor, Dorsomorphin, substantially reduced GLS expression (Figures 3G and S4E), suggesting that GLS is a downstream target of p-AMPK. In ESCC cells with TIGAR overexpression, the ATP levels were significantly lower when there was also an AMPK knockdown while in cells without TIGAR overexpression, the ATP production was not affected by AMPK knockdown (Figure 3H). Accordingly, we observed significant suppression of cancer cell proliferation and colony formation ability when AMPK or GLS expression was knocked down or inhibited, and the suppression effect was much stronger in cells with TIGAR overexpression (Figures 3I–3K and Figures S4F–S4H). Nevertheless, ESCC cell proliferation suppressed by glutamine pathway inhibition could be rescued by α-ketoglutarate (α-KG), a glutamine downstream metabolite that enters the tricarboxylic acid cycle for ATP generation (Figures 3J and 3K; Figures S4G and S4H). In human ESCC tissue samples, we also found strong positive correlations among the protein levels of TIGAR, p-AMPK and GLS measured by IHC (Figure 3L and Table S1).

**Figure 3.**
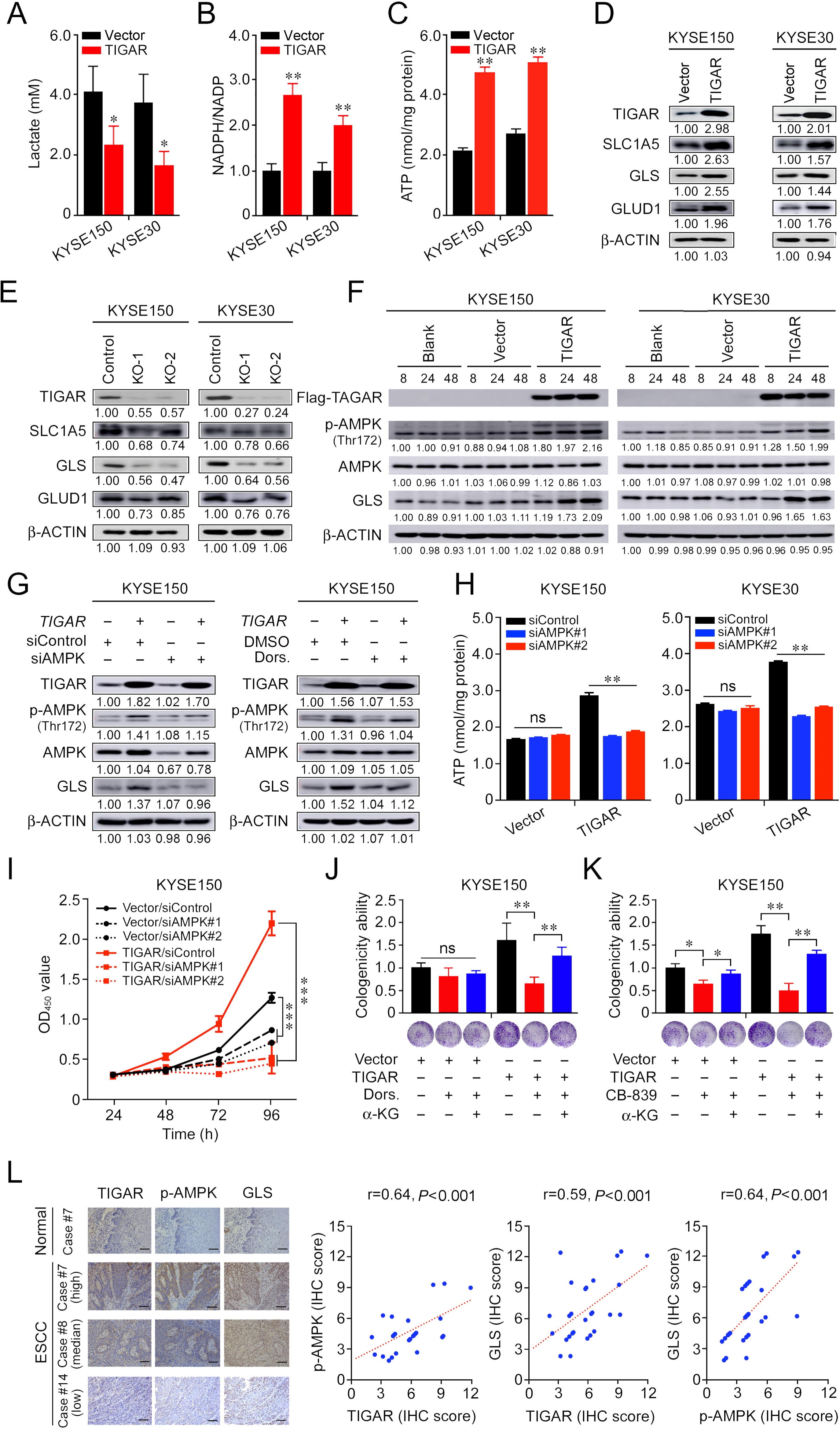
TIGAR inhibits glycolysis but activates glutamine pathway via AMPK to promote ESCC. (A–C) ESCC cells overexpressing TIGAR had significantly reduced intracellular lactate levels (A) but significantly elevated NADPH (B) and ATP levels (C) compared with cells without TIGAR overexpression. (D and E) Western blot analysis of SLC1A5, GLS and GLUD1 protein expression levels in ESCC cells with TIGAR overexpression (D) or TIGAR knockout (E), indicating that glutamine pathway is activated when TIGAR is overexpressed. (F) Western blot analysis of phosphorylated-AMPK (p-AMPK), total AMPK and GLS in ESCC cells overexpressing TIGAR, indicating that both AMPK activation and GLS expression are elevated by TIGAR overexpression. (G) Western blot analysis of GLS in ESCC cells with or without TIGAR overexpression and with or without AMPK knockdown (*left*) or inhibition (*right*). The results indicate that TIGAR promotes the glutamine pathway via activating AMPK. (H) ATP production in ESCC cells with or without TIGAR overexpression by AMPK knockdown, showing that knockdown of AMPK reduces ATP production only in cells with TIGAR overexpression. (I) Knockdown of AMPK significantly inhibited ESCC cell (KYSE150) proliferation and the effect was more pronounced in cells overexpressing TIGAR. (J and K) Inhibitor of AMPK (Dorsomorphin, Dors.) or GLS (CB-839) significantly suppressed ESCC cell colony formation and the effect was more pronounced in cells overexpressing TIGAR. α-Ketoglutarate (α-KG), a metabolite of glutamine, significantly rescued ESCC cell colony formation that was suppressed by AMPK (J) or GLS inhibition (K). (L) The positive correlations among the expression levels of TIGAR, p-AMPK and GLS proteins in human clinical ESCC specimens (n=28). *Left* panel shows the typical pictures of immunohistochemical staining of these 3 proteins in ESCC and normal esophageal tissues. *Right* panel shows the positive correlations among the expression levels of these proteins. Error bars represent SEM obtained from three independent experiments. *, *P*<0.05; **, *P*<0.01 and ***, *P*<0.001 from t-test.

### Glutaminase inhibitor represses primary ESCC and enhances chemotherapy efficacy in mice

We then examined whether glutamine pathway inhibition may suppress the growth and progression of primary ESCC in mice. We intragastically administrated CB-839, a GLS inhibitor, to *Tigar*^+/+^ or *Tigar*^-/-^ mice with 4-NQO induced primary ESCC for 18 days and observed that *Tigar*^+/+^ mice had significantly fewer tumors, smaller tumor size, and a significant recovery from body weight loss as compared with the control group; however, these effects were not observed in *Tigar*^-/-^ mice (Figures 4A–4C and Figure S5A). We examined *Tigar*, p-Ampk and Gls protein levels in mouse ESCC by IHC staining and all three proteins were overexpressed in *Tigar*^+/+^ mice, but not in *Tigar*^-/-^ mice (Figure 4D). Glutamine pathway inhibition by CB-839 also significantly increased the sensitivity of *Tigar*^+/+^ ESCC to routinely used cytotoxic chemotherapeutic agents 5-fluorouracil (5FU) and cisplatin (DDP). We observed that after 3-week treatment with 5FU/DDP, the primary ESCC tumor numbers and size were not significantly reduced compared with those in control mice; however, treatment with combined CB-839 and 5FU/DDP remedy significantly reduced ESCC numbers and size (Figures 4E and 4F; Figure S5B) and prolonged animal survival compared with treatment of 5FU/DDP alone (Figure 4G). Immunohistochemistry staining showed substantially lower Ki67 and higher cleaved CASPASE-3 proteins in mouse ESCC treated with combined remedy compared with that in mouse ESCC treated with 5FU/DDP or CB-839 alone (Figure 4H). These results clearly demonstrate that glutamine pathway inhibition has the synergic effect with cytotoxic chemotherapeutic agents on repressing *Tigar*-overexpressing primary ESCC in mice.

**Figure 4.**
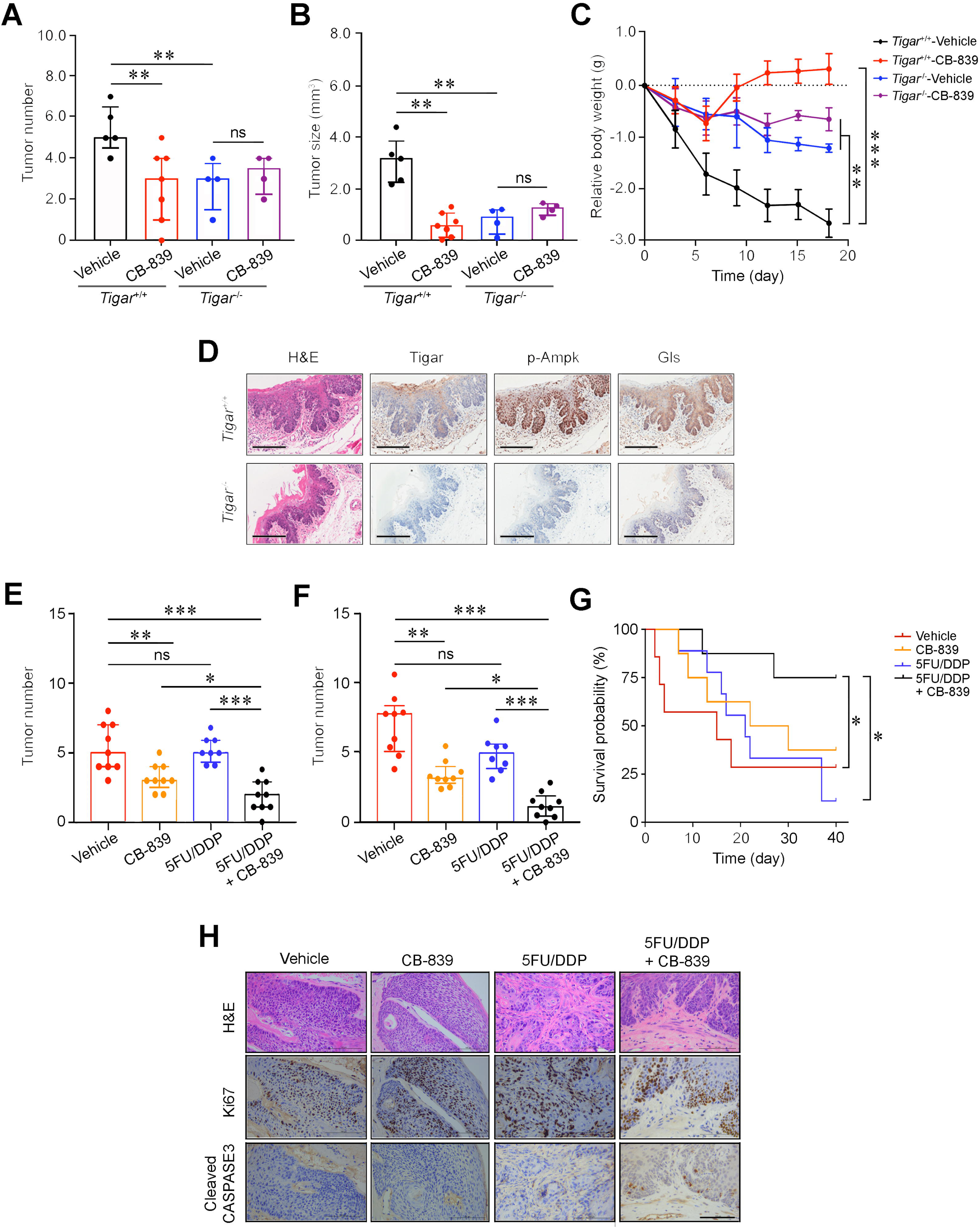
Glutaminase inhibitor represses primary ESCC and enhances chemotherapy efficacy in mice. (A and B) Treatment with CB-839 significantly reduced primary ESCC tumor number (A) and tumor size (B) induced by 4-NQO in mice with *Tigar*^+/+^ but not in mice with *Tigar*^-/-^. Mice were given 4-NQO (see Methods for details) for 16 weeks and after 28 weeks they received CB-839 (200 mg/kg, p.o.) treatment for 18 days. **, *P*<0.01 from Mann-Whitney test. (C) Comparison of body weight gain among *Tigar*^+/+^ or *Tigar*^-/-^ mice bearing ESCC with or without CB-839 treatment. **, *P*<0.01 and ***, *P*<0.001 from Mann-Whitney test. (D) Immunohistochemical staining of *Tigar*, p-Ampk, Gls in mice ESCC induced by 4-NQO, showing that the Ampk activation and Gls expression were only seen in *Tigar*^+/+^mice but not in *Tigar*^-/-^. (E and F) Treatment with CB-839 (200 mg/kg, p. o., every day) or 5FU (20 mg/kg, i. p., every 3 days)/DDP (2 mg/kg, i. p., every 5 days) separately or in their combination as indicated in the Figure labels in by 4-NQO-induced mice with *Tigar*^+/+^. Combined treatment of 5FU/DDP and CB-839 significantly reduced ESCC tumor number (E) and tumor size (F). *, *P*<0.05; **, *P*<0.01 and ***, *P*<0.001 from Mann-Whitney test. (G) Combined treatment of 5FU/DDP and CB-839 significantly prolonged survival time of mice with ESCC induced by 4-NQO with *Tigar*^+/+^. *, *P*<0.05 from log-rank test. (H) Hematoxylin and eosin (H&E) staining (*top*) and immunohistochemical staining of Ki67 and cleaved CASPASE 3 (*middle* and *bottom*) in 4-NQO-induced primary ESCC n mice with *Tigar*^+/+^, showing that combined treatment of 5FU/DDP and CB-839 inhibited proliferation and promoted apoptosis of cancer cells.

### Glutamine pathway is a therapeutic target for TIGAR-overexpressing human ESCC

In vitro assays showed that glutamine deprivation or GLS expression knockdown significantly inhibited KYSE150 and KYSE30 proliferation and the inhibitory extent was much greater in TIGAR-overexpressing cells than control cells (Figures S6A and S6B). The ATP levels were remarkably declined in ESCC cells deprived of glutamine or with GLS knockdown and this result seemed to occur only in cells with but not without TIGAR overexpression (Figures S6C and S6D). Based on these results, we treated KYSE150 and KYSE30 with CB-839 or 5FU/DDP and found that while both treatments significantly suppressed ESCC cell proliferation in dose-dependent manner, ESCC cells with TIGAR overexpression seemed more vulnerable to CB-839 but more resistant to 5FU/DDP remedy than ESCC cells without TIGAR overexpression (Figures 5A and 5B; Figures S6E and S6F). Treatment of tumor xenografts derived from TIGAR-overexpressing ESCC cells in mice showed similar results: glutamine deprivation or CB-839 treatment alone significantly suppressed tumor growth by more than 50% while 5FU/DPP alone had no significant effects under our experimental conditions; however, combination of glutamine deprivation or CB-839 with 5FU/DDP significantly repressed tumor xenograft growth (Figures 5C and 5D). Next, we established patient-derived xenograft (PDX) mouse models to investigate whether glutamine pathway inhibition has the effect on repressing TIGAR-overexpressing human ESCC. TIGAR overexpression and glutamine pathway activation in primary ESCC of patients and PDXs of mice were determined and verified by immunohistochemical staining (Figures S7A–S7D). In mice, PDXs with high TIGAR expression grew faster than PDXs with low TIGAR expression (Figures 5E and 5F; Figures S7E-S7J), consistent with the aforementioned results showing that TIGAR plays an important part in ESCC progression. Based on the results shown above, mice carrying PDX with TIGAR overexpression were treated with only 5FU/DDP or 5FU/DDP plus CB-839 and mice carrying PDX without TIGAR overexpression were treated with only 5FU/DDP. We found that PDXs with high TIGAR expression were more resistant to 5FU/DPP treatment than PDX with low TIGAR expression. However, treatment with combined CB-839 and 5FU/DDP significantly repressed the growth of PDXs (Figures 5E and 5G; Figures S7E **and** S7H), which was characterized by decreased proliferation marker Ki67 and increased apoptosis marker cleaved CASPASE-3 levels (Figures 5G and 5H; Figures S7K and S7L).

**Figure 5.**
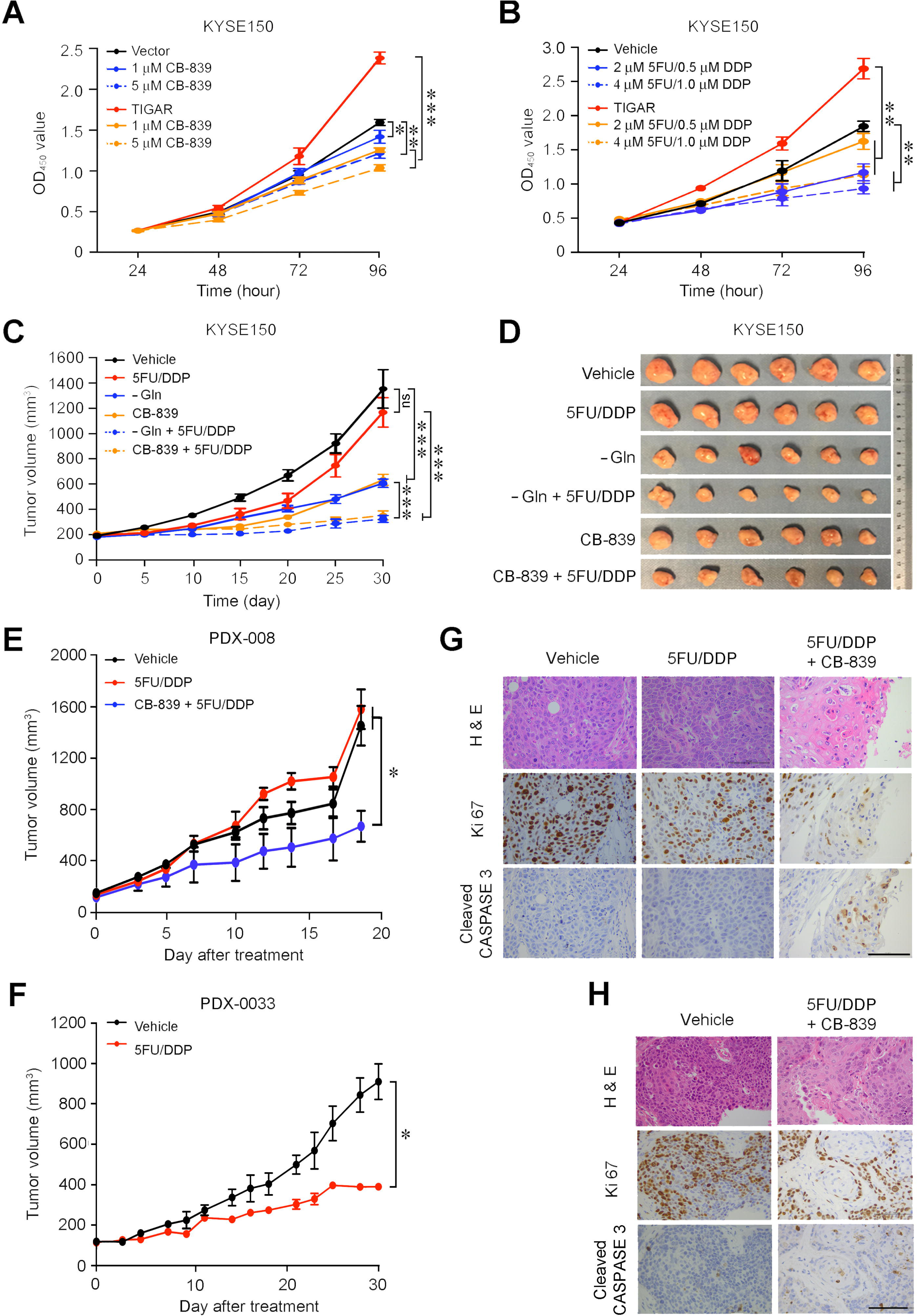
Glutamine pathway is a therapeutic target for TIGAR-overexpressing human ESCC. (A) Dose-dependent inhibition of ESCC cell (KYSE150) proliferation by GLS inhibitor CB-839. The effect was more pronounced in cells overexpressing TIGAR. Error bars represent SEM obtained from three independent experiments. *, *P*<0.05; **, *P*<0.01 and ***, *P*<0.001 from Student’s *t*-test. (B) Dose-dependent inhibitory effect of 5FU/DDP on ESCC cell (KYSE150) proliferation. The efficacy was less pronounced in cells overexpressing TIGAR, suggesting chemoresistance. Error bars represent SEM obtained from three independent experiments. **, *P*<0.01 from Student’s *t*-test. (C and D) Treatment of mice transplanted ESCC with 5FU (20 mg/kg, i. p., every 3 days)/DDP (2 mg/kg, i. p., every 5 days), Gln deprivation or CB-839 (200 mg/kg, p.o., every day) separately or in combination as indicated in the Figure labels. Combined treatment of Gln deprivation or CB-839 with 5FU/DDP significantly repressed xenograft tumor growth as shown in tumor growth curves (C) and tumor size in the experiment end at 21 days (D). ***, *P*<0.001 from Mann-Whitney test. (E) Combined treatment of 5FU/DDP with CB-839 significantly repressed the growth of patient derived xenograft (PDX) with high TIGAR expression in mice *, *P*<0.05 from Student’s *t*-test. (F) 5FU/DDP treatment alone significantly repressed the growth of PDX with low TIGAR expression in mice. *, *P*<0.05 from Student’s *t*-test, suggesting more sensitive to the chemotherapy compared with PDX with high TIGAR expression. (G) Hematoxylin and eosin (H&E) staining (*top*) and immunohistochemical staining of Ki67 and cleaved CASPASE 3 (*middle* and *bottom*) in PDX with high TIGAR expression, showing that combination treatment of 5FU/DDP with CB-839 suppressed proliferation and promoted apoptosis of PDX cancer cells. (H) Hematoxylin and eosin (H&E) staining (*top*) and immunohistochemical staining of Ki67 and cleaved CASPASE 3 (*bottom*) in PDX with low TIGAR expression, showing that 5FU/DDP treatment inhibited proliferation and promoted apoptosis of PDX cancer cells.

## Discussion

Our previous study identified *TIGAR*, among others, as a copy-number gain gene in ESCC (Chang et al., 2017). In the present study, we have demonstrated that TIGAR is overexpressed in the majority of human ESCC, confirming previous RNA-seq and mRNA microarray results (Chang et al., 2017; Su et al., 2011). Additionally, we have found that high TIGAR expression levels are significantly correlated with advanced tumor stages, lymph node metastasis and poor patient survival. Animal experiments also demonstrate that *Tigar* plays an important role in chemical carcinogen-induced ESCC carcinogenesis. In an effort to explain the underlying mechanism, we have revealed that aberrant TIGAR expression inhibits glycolysis but activates the glutamine pathway, which may facilitate ESCC cell survival and progression. More importantly, based on these results, we have performed treatment targeting glutamine pathway in mouse primary ESCC and PDX models and found that inhibition of glutamine pathway significantly repress ESCC progression and enhance the efficacy of cytotoxic chemotherapy (5FU/DDP). Together, these results demonstrate that TIGAR overexpression is essential for the development and progression of ESCC through a underlying mechanism of glutamine pathway activation; thus, inhibition of aberrant TIGAR expression dependent glutamine pathway may be a therapeutic target for precision treatment of this malignancy (Figure 6).

**Figure 6.**
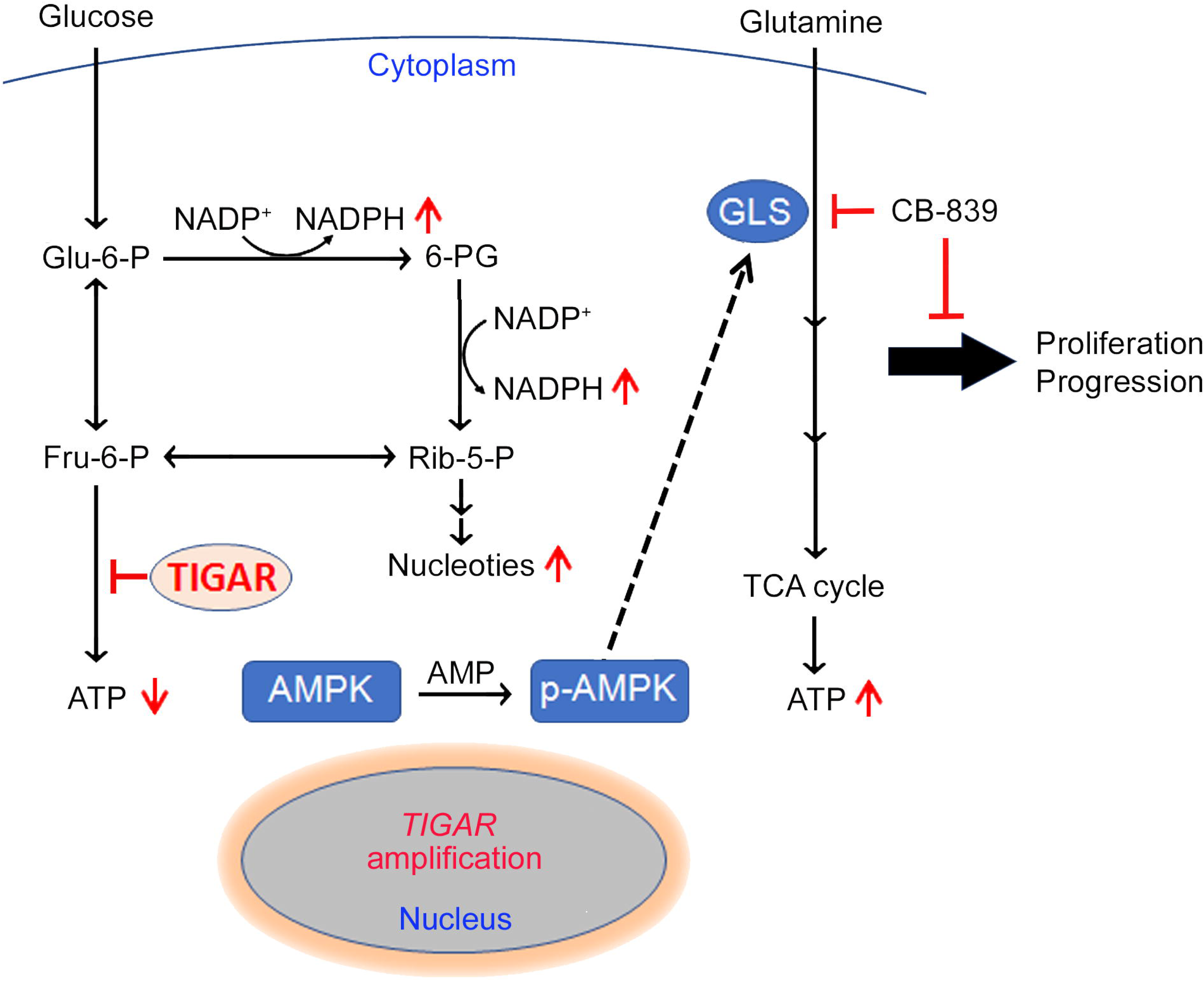
A proposed model for the role of TIGAR overexpression and the effect of inhibition of glutamine pathway in ESCC progression. TIGAR overexpression inhibits glycolysis but remodels glutamine pathway, which produces more anti-oxidants to protect cancer cells from reactive oxygen species killing and more energy to promote cancer cell proliferation and progression. Inhibiting glutamine pathway therefore might suppress the progression of ESCC with TIGAR overexpression.

We detected a significantly reduced lactate level but a significantly increased NADPH level in ESCC overexpressing TIGAR, which is consistent with the role of TIGAR in glucose metabolism and the proposed consequences of its overexpression. Since glycolysis is the major energy source for cancer cells, one would expect that inhibition of glycolysis may bring down the ATP levels in ESCC cells with TIGAR overexpression; however, we observed a significantly elevated rather than decreased ATP level in these cells. This paradoxical result encouraged us to explore the other energy providers for ESCC cell survival and proliferation and led to uncover the glutamine pathway activation in ESCC cells overexpressing TIGAR. Metabolic remodeling such as glutamine addiction is common in many types of human cancer and has been associated with malignant phenotypes (DeBerardinis and Chandel, 2016; Hanahan and Weinberg, 2011; Martinez-Outschoorn et al., 2017), but the underlying causes remain poorly understood. Here, we demonstrate for the first time that TIGAR overexpression in ESCC cells can drive this metabolic remodeling and facilitates the tumor progression.

In this study, we have shown that the transient ectopic overexpression of TIGAR in ESCC cells can induce phosphorylation of AMPK, a well-known central regulator of cellular response to energetic stress (Garcia and Shaw, 2017), in a time-dependent manner. We observed that the expression level of GLS, a rate-limiting enzyme in the glutamine pathway, is concurrently increased when AMPK is activated by TIGAR. Conversely, when AMPK activity is knocked down or inhibited, the GLS expression level is also concurrently decreased. We also found a significantly positive correlation among TIGAR, p-AMPK and GLS protein levels in human ESCC samples and in carcinogen-induced *Tigar*^+/+^ mice ESCC. These results clearly indicate that TIGAR-overexpressing ESCC cells restore cellular energy homeostasis by using the glutamine pathway via activating AMPK. Consistent with our findings, the activation of AMPK has been shown to promote the glutamine pathway for ATP renewal in T cells (Blagih et al., 2015). However, how p-AMPK activates the glutamine pathway remains to be elucidated. In particular, it would be interesting to investigate whether GLS is a direct p-AMPK substrate or its expression is indirectly induced by p-AMPK signaling.

Genomic alteration caused metabolic remodeling could make cancer cells addicted to certain nutrients, which can then become a potential therapeutic target. In the present study, we found that GLS inhibitor CB-839 can significantly repress human ESCC cell-derived xenografts, chemical carcinogen-induced primary ESCC and PDXs in mice and the effect is more pronounced and sometimes only observed in tumors overexpressing TIGAR. Furthermore, in these models, we found that ESCC cells overexpressing TIGAR are more resistant to 5FU/DDP, the first-line chemotherapeutic agents routinely used to treat advanced or recurrent ESCC. This could be due to the other metabolic pathways (e. g., glutamine and pentose phosphate) that cancer cells shift to since TIGAR overexpression produces more antioxidants against apoptosis and more intermediates for DNA synthesis (Lee et al., 2014; Wanka et al., 2012). Interestingly, we found that inhibition of glutamine pathway enhances the sensitivity of TIGAR overexpressing ESCC cells towards cytotoxic anticancer agents, 5FU/DDP, suggesting that a combination of glutamine pathway inhibitors and conventional cytotoxic agents may be a promising novel treatment option for ESCC with TIGAR overexpression. This finding is particularly important because no target therapy or specifically effective anticancer drugs are currently used for ESCC.

Our present study also contributes to the existing literature by identifying the activation of the glutamine pathway as a consequence of TIGAR overexpression in ESCC. Unfortunately, how TIGAR expression is regulated is still poorly understood. TIGAR expression regulation was initially identified through microarray analysis of gene expression following TP53 activation (Bensaad et al., 2006; Jen and Cheung, 2005); but several later studies have shown that TIGAR expression is not all dependent on TP53 (Cheung et al., 2013; Pena-Rico et al., 2011). It has also been reported that c-MET inhibition may result in marked downregulation of TIGAR and intracellular production of NADPH in nasopharyngeal cancer (Lui et al., 2011). In the present study, we found the TIGAR overexpression in ESCC is not correlated with *P53* or *c-MET* expression (Figures S8A and S8B), but is associated with somatic copy number gain of the gene itself (Figure S8C). The accelerated growth of cancer could make the tumor genome subject to even more alterations. Since TIGAR amplification and overexpression have been found in many types of human primary cancer based on our analysis on TCGA data (Figure S9), our results in the current study may extend to other cancer types where TIGAR is overexpressed.

In summary, the present study demonstrates that the frequently amplified TIGAR gene is overexpressed in the majority of human ESCC and plays an important oncogenic role in ESCC progression. One of the underlying mechanisms is that TIGAR overexpression inhibits glycolysis to produce more antioxidants for cancer cell survival and, on the other hand, TIGAR reprograms ESCC cell metabolism using glutamine to produces more energy for cancer progression. Based on these findings, we suggest that the aberrant TIGAR overexpression may sever as a biomarker for precision ESCC treatment and inhibition of TIGAR-dependent glutamine pathway activation is a promising target therapy for ESCC. These findings might also provide the rationale for clinical trials testing glutamine pathway inhibitors in combination with chemotherapy in TIGAR-expressing ESCC.

## STAR METHODS

### KEY RESOURCES TABLE

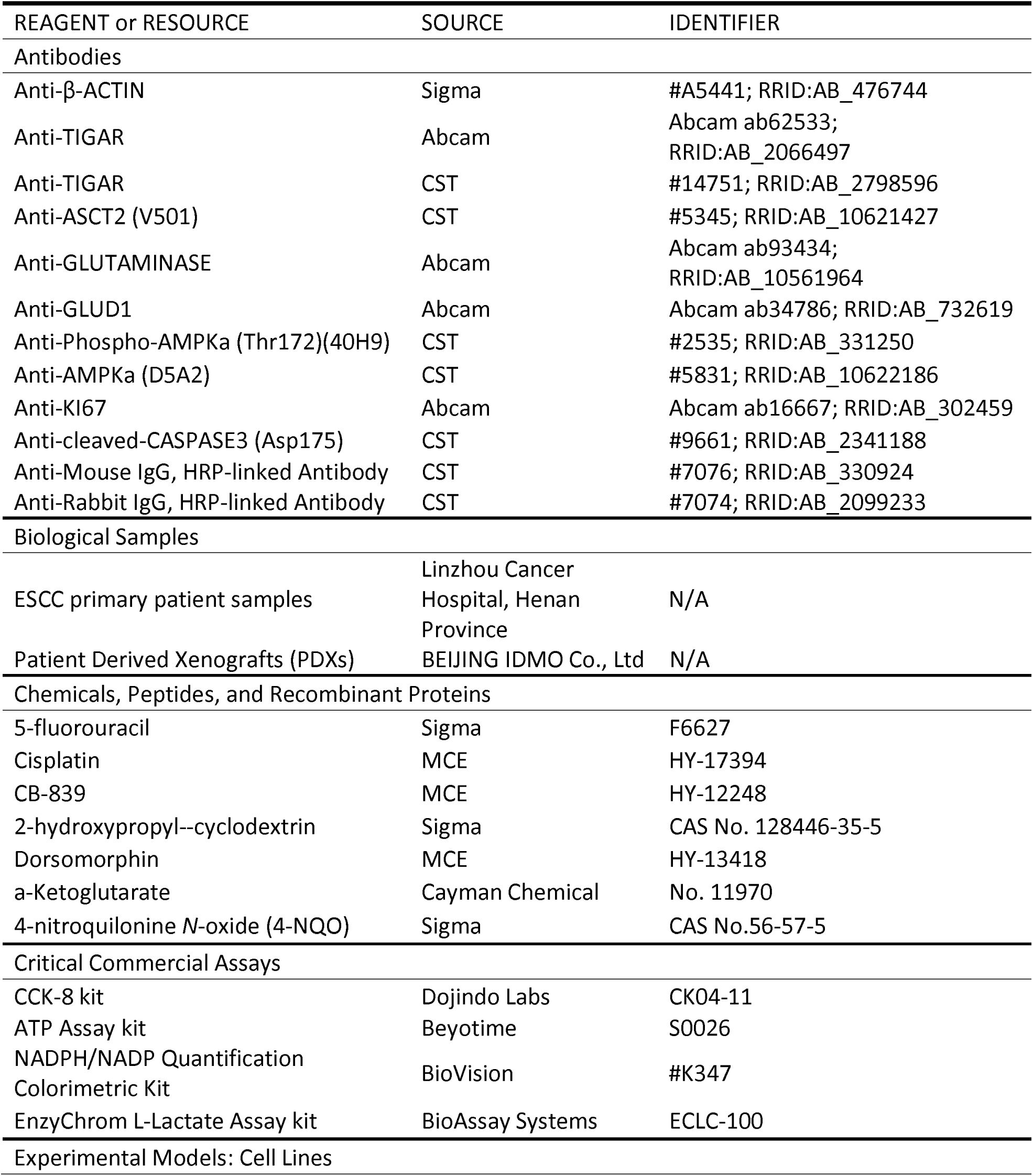
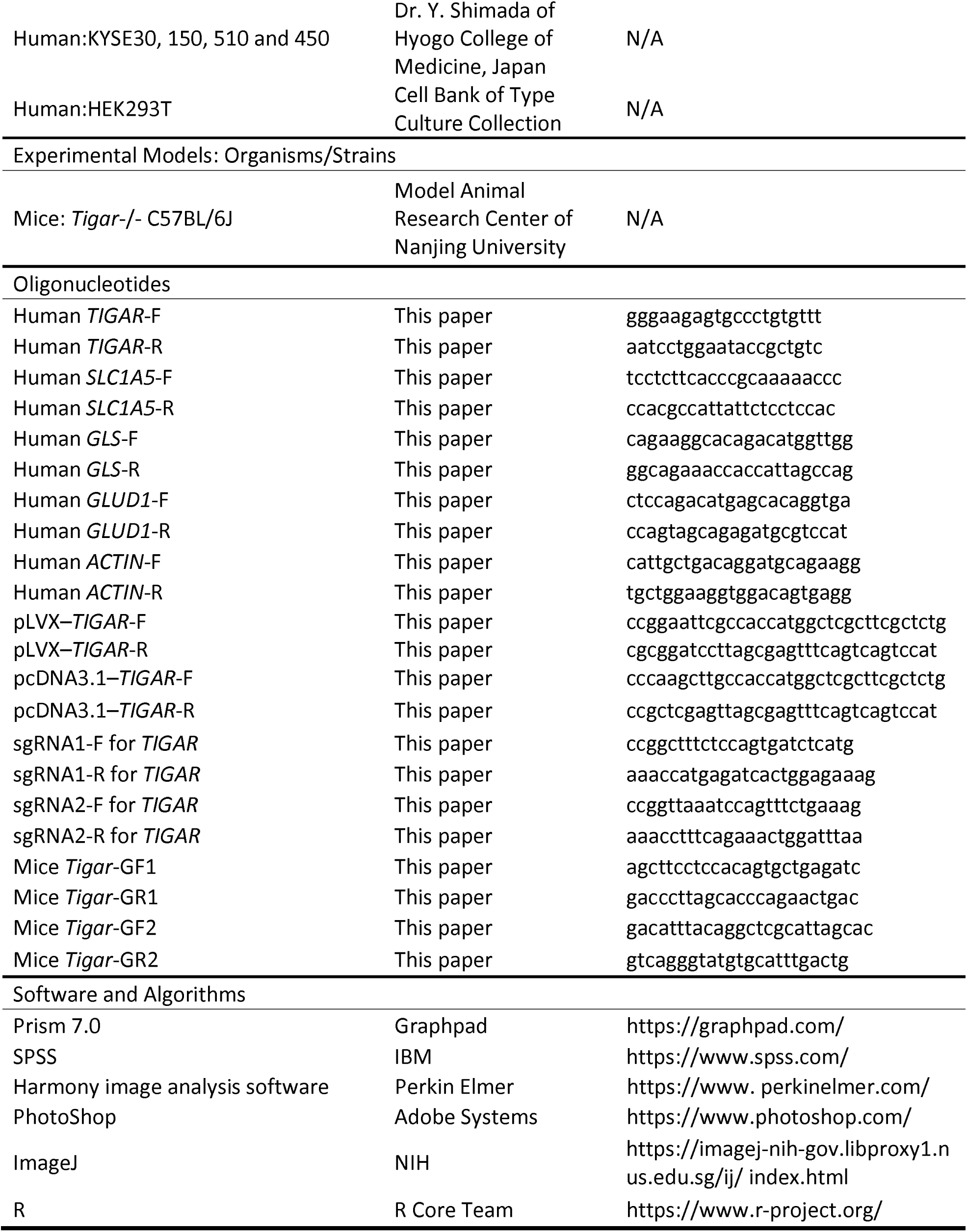

### LEAD CONTACT AND MATERIALS AVAILABILITY

Further information and requests for resources and reagents should be directed to and will be fulfilled by the Lead Contact, Jiahui Chu or Xiangjie Niu

### EXPERIMENTAL MODEL AND SUBJECT DETAILS

#### Cell Lines and Cell Culture

Human ESCC cell lines KYSE30, KYSE150, KYSE450 and KYSE510 were kind gifts from Dr. Y. Shimada of Hyogo College of Medicine, Japan. HEK293T cells were purchased from the Cell Bank of Type Culture Collection of Chinese Academy of Sciences, Shanghai Institute of Biochemistry and Cell Biology. All cell lines were authenticated by DNA finger printing analysis, confirmed to be free of mycoplasma infection and cultured respectively in DMEM (HEK293T) or RPMI 1640 (all other cell lines) medium (Hyclone) supplemented with 10% fetal bovine serum at 37°C in a humidified incubator with 5% CO_2_.

#### Study subjects and tissue specimens

This study enrolled a panel of 225 patients with ESCC from Linzhou Cancer Hospital, Henan Province between 2015 and 2016. All patients underwent esophagectomy without chemotherapy or radiotherapy before surgery. Surgically removed tumor and their corresponding adjacent normal esophageal tissue (≥5 cm from tumor site) samples were collected from each patient at the time of surgery. All tumors were histopathologically classified as ESCC. Relevant characteristics and clinical information were obtained from patients’ medical records (Table S2). Informed consents were obtained from all individuals and this study was approved by Chinese Academy of Medical Sciences Cancer Hospital.

#### Animals

BALB/c nude mice (4-6 week old) were purchased from Huafukang Biotechnology, Beijing. 6-8 weeks old C57BL/6 mice and *Tigar*^-/-^ C57BL/6J mice purchased from Model Animal Research Center of Nanjing University. Mice were maintained in individual ventilated cages and fed with autoclaved food and water in the Animal Care Center, Chinese Academy of Medical Sciences. All animal experiments were carried out in compliance with approved protocols and guidelines from the Institutional Animal Care and Use Committee of the Chinese Academy of Medical Sciences. Kaplan-Meier survival curves were compared using the Mantel-Cox Log-rank test via Graphpad Prism 7.

### METHOD DETAILS

#### Selection of candidate genes for potential functional screening

We previously identified 23 focal amplified regions overlapping 1,591 genes (Chang et al., 2017). Of them, 149 genes were significantly overexpressed in tumor samples (differential expression analysis P<0.05, fold change >1.35) and their expression levels were significantly correlated with their copy-number gains (Spearman’s correlation coefficient >0.35, P<0.05). These 149 genes (Figure S3) were chosen for functional screening in the present study.

#### RNA interfering-based high content screening assays

The small interfering RNA (siRNA) library provided by Dharmacon comprised 3 individual non-overlapping siRNA designs for each gene and the repression efficiency was guaranteed by the provider. The sequences specific to the candidate genes are shown in Table S3. The high content screening assays were performed as described previously (McCoy, 2011; Zanella et al., 2010). Briefly, cells were reverse transfected with siRNAs in 96-well plates using Lipofectamine® RNAiMAX Transfection Reagent (Life Technologies) following the manufacturer’s instructions. Ten μl of siRNA (25 nM) solution and 10 μl of transfection mix were placed in the plates and after incubation at room temperature for 20 min, about 3,000 cells in 80 μl of 1640 medium were seeded per well and incubated for 3 days at 37°C. Cells were then fixed and permeabilized with 5% paraformaldehyde (Sigma) and 0.2% Triton X-100 (Sigma) for 45 min. To prevent non-specific binding, cells were incubated with 3% bovine serum albumin (Gerbu) and 0.05% Triton X-100 for 30 min. Nuclei and Actin were then stained with 100 ng/ml DAPI (C1002, Beyotime) and 67 ng/ml phalloidin labeled with tetra-methylrhodamine isothiocyanate (Sigma) in blocking solution at 4°C overnight. After washing with PBS, fluorescence images of cells were acquired with an Image Analyzer (Perkin Elmer). Nuclei were segmented by adaptive thresholding and the number of segmented nuclei was used as a proxy for cell count. Baseline and main effects were computed from non-targeting controls and single-gene knockdowns for each siRNA design. P values were computed by a t-test over 3 replicates for each cell. *EGFR*, a known driver gene that promotes ESCC cell proliferation, was used as a positive control for the screening assays.

#### RNA preparation and quantitative real-time PCR analysis

Total RNA from cells or tissue specimens was isolated by using TRIzol Reagent (Invitrogen) and converted to cDNA with the PrimeScriptTM RT reagent kit (TaKaRa). Quantitative real-time PCR (qRT-PCR) was accomplished in triplicate using the SYBR-Green method on an ABI 7900HT Real-Time PCR system. Individual RNA expression level was determined relative to the level of β-*ACTIN* RNA. The primer sequences used for PCR are shown in Key resources table.

#### Western blot analysis

Total proteins extracted from tissue samples or cell lines were subjected to SDS-PAGE and transferred to PVDF membranes (Millipore). Antibody against TIGAR (ab62533), GLS (ab93434) or GLUD1 (ab34786) from Abcam, antibody against phosphorylated AMPKα at Thr172 (2535), AMPKα (5831) or ASCT2 (SLC1A5; 3545) from CST and antibody against β-ACTIN (sc-47778) from Santa-Cruz were used. The membranes were incubated with the primary antibody and visualized with a Phototope-Horseradish Peroxidase Western Blot detection kit (Cell Signaling Technology). The protein bands were quantified by gray scanning.

#### Plasmids and lentiviral production as well as transduction

Full length of human TIGAR cDNA with artificial BamH I and EcoR I enzyme restriction sites was PCR-amplified and subcloned to the lentiviral expression vector pLVX-IRES-Neo, which was then transfected into HEK293T cells to produce viruses. Lentiviral supernatant was harvested at 48 or 72 hours post-transfection. KYSE150 and KYSE30 cells were infected with concentrated viruses and cultured with complete medium for 24 hours followed by selection with G418. To construct expression vectors of Flag-tagged TIGAR, cDNA encoding TIGAR was subcloned to pcDNA3.1-Flag, yielding pcDNA3.1-Flag-TIGAR (Key resources table).

#### Establishment of TIGAR-knockout cell lines by CRISPR editing

The CRISPR/Cas9 system was used to generate genomic deletion of *TIGAR* in ESCC cell lines. Single-guide RNA (sgRNA) sequences designed to target the genomic sequence of TIGAR were cloned into plasmid pUC19-U6-sgRNA. The pCAG-Cas9-EGFP and pUC19-U6-sgRNA plasmids were co-transfected into HEK293T cells and the fluorescent cells were sorted via flow cytometry. DNA was extracted from harvested cells and the target fragment was amplified and PCR products were re-annealed to generate hetero-duplexed DNA. Then T7EI assay were carried out to confirm the editing efficiency (Koike-Yusa et al., 2014; Zhou et al., 2014). Two sgRNAs with high efficiency were selected and cloned into plasmid PB-U6-Bbs1-sgRNA-Neo (Key resources table). KYSE150 and KYSE30 cells were co-transfected with PB-U6-Bbs1-sgRNA-Neo and PBase and cultured with complete medium for 24 hours followed by selection with G418.

#### Cell viability and colony formation assays

Cell viability was measured using the CCK-8 kit (Dojindo Labs). Colony formation ability was determined by counting the number of cells in 12-well cell-culture cluster with complete growth medium after fixing with methanol and staining with crystal violet.

#### Measurement of ATP, NADP, NADPH and lactate levels

The intercellular ATP, NADP, NADPH and extracellular lactate levels were measured using the ATP Assay kit (Beyotime), the NADPH/NADP Quantification Colorimetric Kit (BioVision, #K347), or the EnzyChrom L-Lactate Assay kit (BioAssay Systems), respectively, as described by the manufacturers.

#### In vitro invasion and migration assays

Invasion assay was done in a 24-well Millicell chamber. The 8-μm pore inserts were coated with Matrigel (BD Biosciences). Cells (7×104) were added to coated filters in serum-free medium in triplicate wells. RPMI 1640 media containing 20% FBS was added to the lower chamber as chemo-attractant. After 24 h at 37°C in a 5% CO2 incubator, the Matrigel coating on the upper surface of the filter was removed. Cells that migrated through the filters were fixed with methanol, stained with 0.5% crystal violet, and photographed. Cell number on 3 random fields was counted. The migration assay was conducted in a similar fashion without coating with Matrigel.

#### Xenograft tumor formation and anticancer treatment

To examine xenograft tumor formation ability of various ESCC cells, female BALB/c nude mice (5 animals in each group) were subcutaneously injected with ESCC cells (1×10^6^) and raised for 6 weeks. Tumor volume was measured every week and defined by length x width^2^ × 0.5. For anticancer treatment assays, xenograft tumor models were established by subcutaneously inoculating cancer cells into the armpit of one animal. Four weeks later, the tumor was isolated and cut into pieces (1.5 mm thick) and transplanted into armpit of the other mice. When tumor size reached approximate 200 mm^3^, mice were randomly divided into groups (n=6 per group) and treated with 5FU (i. p.), DDP (i. p.), CB-839 (p. o.), glutamine deprivation, or their combination. Mice treated with 2-hydroxypropyl-20-cyclodextrin were served as vehicle control. The doses administrated are detailed in the corresponding figure legends. The glutamine deficient diet was purchased from Jiangsu Synergy Laboratory Animal Feed Supplies. Animal experiments were carried out in compliance with approved protocols and guidelines from the Institutional Animal Care and Use Committee of the Chinese Academy of Medical Sciences.

#### Induction of ESCC by 4-NQO in genetically *Tigar*-engineered mice and anticancer treatment

We used chemical carcinogen 4-NQO to induce ESCC in genetically engineered *Tigar*^+/+^, *Tigar*^+/-^ or *Tigar*^-/-^ C57BL/6J mice purchased from Model Animal Research Center of Nanjing University. The animal *Tigar* genotypes were verified by PCR with the primers shown in Key resources table. Mice (n=11 per group) were treated with 4-NQO (100 μg/ml) or 2% propylene glycol (vehicle control) in drinking water for 16 weeks and then followed up for another 12 weeks after 4-NQO withdrawal (Tang et al., 2004). The stock solution of 4-NQO in propylene glycol (5 mg/ml) was weekly prepared and 1:50 diluted in drinking water for use, which was changed once a week. Mice were allowed access ad *libitum* to the drinking water at all times during the treatment. For CB-839 treatment, both *Tigar*^+/+^ and *Tigar*^-/-^ mice were received 4-NQO as aforementioned and 28 weeks later, they were intragastrically treated with CB-839 (200 mg/kg body weight) or 25% 2-hydroxypropyl-21-cyclodextrin as vehicle control once a day for 4 weeks. Mice were scarified in the end of the experiments and the esophagus was dissected longitudinally and tumors at diameter of ≥0.5 mm were counted and the length and width of tumor were measured to calculate tumor volume (length × wideth^2^ × 0.5). All animals were weighted every three days. The esophagus was fixed in buffered formalin and embedded in paraffin. Sections of the paraffin-embedded tissues were used for ESCC diagnosis and for immunohistochemical (IHC) examination. The differences of survival time among *Tigar*^+/+^, *Tigar*^+/-^ and *Tigar*^-/-^ mice were compared by Kaplan-Meier plotting.

#### Establishment of mice PDX models and anticancer treatment

Surgically removed fresh human ESCC tumor sample (F0 tumors) was implanted subcutaneously in NOD/SCID/IL-2Rγnull (NSG) mice for PDX expanding, which was designated as F1 tumor. If necessary, F2 tumor was made in another mouse using F1 tumor. When expending to a size of about 800 mm^2^, F1 or F2 tumor was removed from animal and cut into small pieces and implanted again in a group of 5 or 6 mice. Histopathology and IHC of TIGAR, p-AMPK and GLS proteins in PDX were analyzed and compared with F0 tumor. When PDX reached 100-200 mm^3^, mice were randomly divided into groups for anticancer treatment. Mice carrying PDX with high TIGAR overexpression received 5FU/DDP or CB-839 plus 5FU/DDP while mice carrying PDX without TIGAR overexpression received 5FU/DDP only. 5FU was given twice a week at a dose of 20 mg/kg body weight and DDP was given once every 5 days at 2 mg/kg body weight. CB-839 was administrated very day at 200 mg/kg body weight. Control mice were given corresponding solvent. Tumor size was examined every 3 days by caliper measurements. Animals were sacrificed when the tumors in vehicle control group reached 2,000 mm^3^.

#### Immunohistochemical analysis

Paraffin-embedded tissue sections and tissue arrays were used for immunohistochemical examination of TIGAR, p-AMPK, GLS, Ki-67 and cleaved CASPASE3 expression levels. Briefly, the sections were incubated with primary antibody against TIGAR (1:200; Abcam, ab62533), p-AMPK (1:100; CST, 2535), GLS (1:100; Abcam, ab93434), Ki-67 (1:100; Abcam, ab16667) or cleaved CASPASE3 (1:200, CST, #9661) at 4°C overnight and then detected with the ABC Kit (Pierce). The labeling score of intensity was estimated as negative (0), weak (1), moderate (2) and strong (3). The extent of staining, defined as the percentage of positive stained cells, was scored as 1 (≤10%), 2 (11%–50%), 3 (51%–80%) and 4 (>80%). The total immune reactive score (IRS) was obtained by multiplying the score of intensity and that of extent, ranking from negative (–) to >6 (+++).

#### Data mining

The cBioPortal for cancer genomics (http://cbioportal.org) was used to analyze copy number variations of TIGAR in various types of cancer (date updated in May 2018). The Oncomine database (http://www.omcomine.org) was queried to see if TIGAR overexpression exists across previous ESCC studies.

#### Statistical analysis

We calculate Spearman’s correlations between copy number and mRNA expression level of specific genes and the correlation is deemed significant and positive when r>0.35 and (nominal) *P*<0.05. Functional analysis results were presented as mean ± SEM of three or more experiments. The Student’s t-test was used to compare two groups whenever the data showed a normal distribution; otherwise, the Mann-Whitney U test was used. *P*<0.05 was considered significant for all statistical analyses.

## Supporting information

Supplementary Figue S1

Supplementary Figue S2

Supplementary Figue S3

Supplementary Figue S4

Supplementary Figue S5

Supplementary Figue S6

Supplementary Figue S7

Supplementary Figue S8

Supplementary Figue S9

