## Supplementary figures and images for "Metabolic remodeling by TIGAR overexpression is a therapeutic target in esophageal squamous-cell carcinoma"

### Supplementary Figue S1

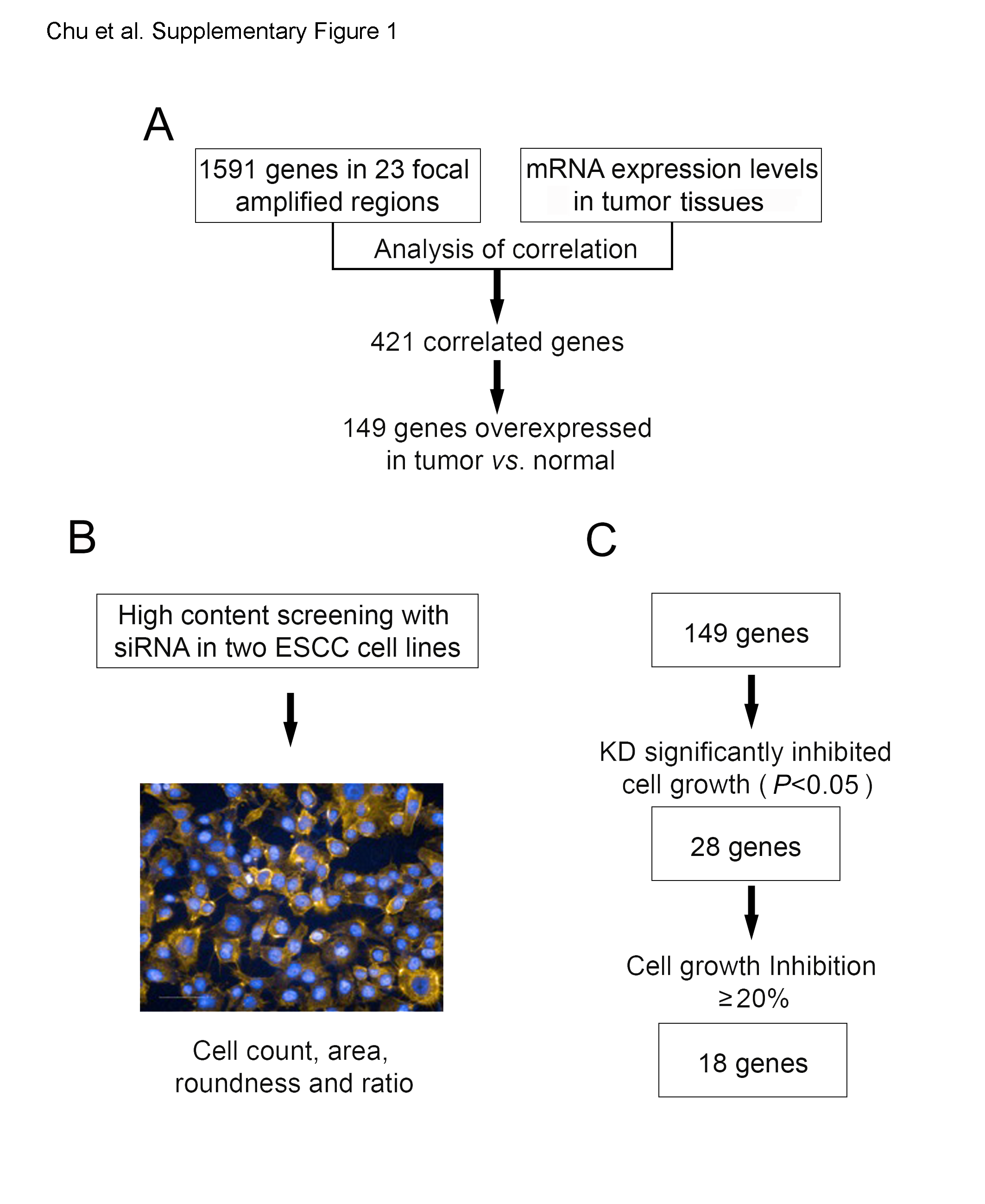

### Supplementary Figue S2

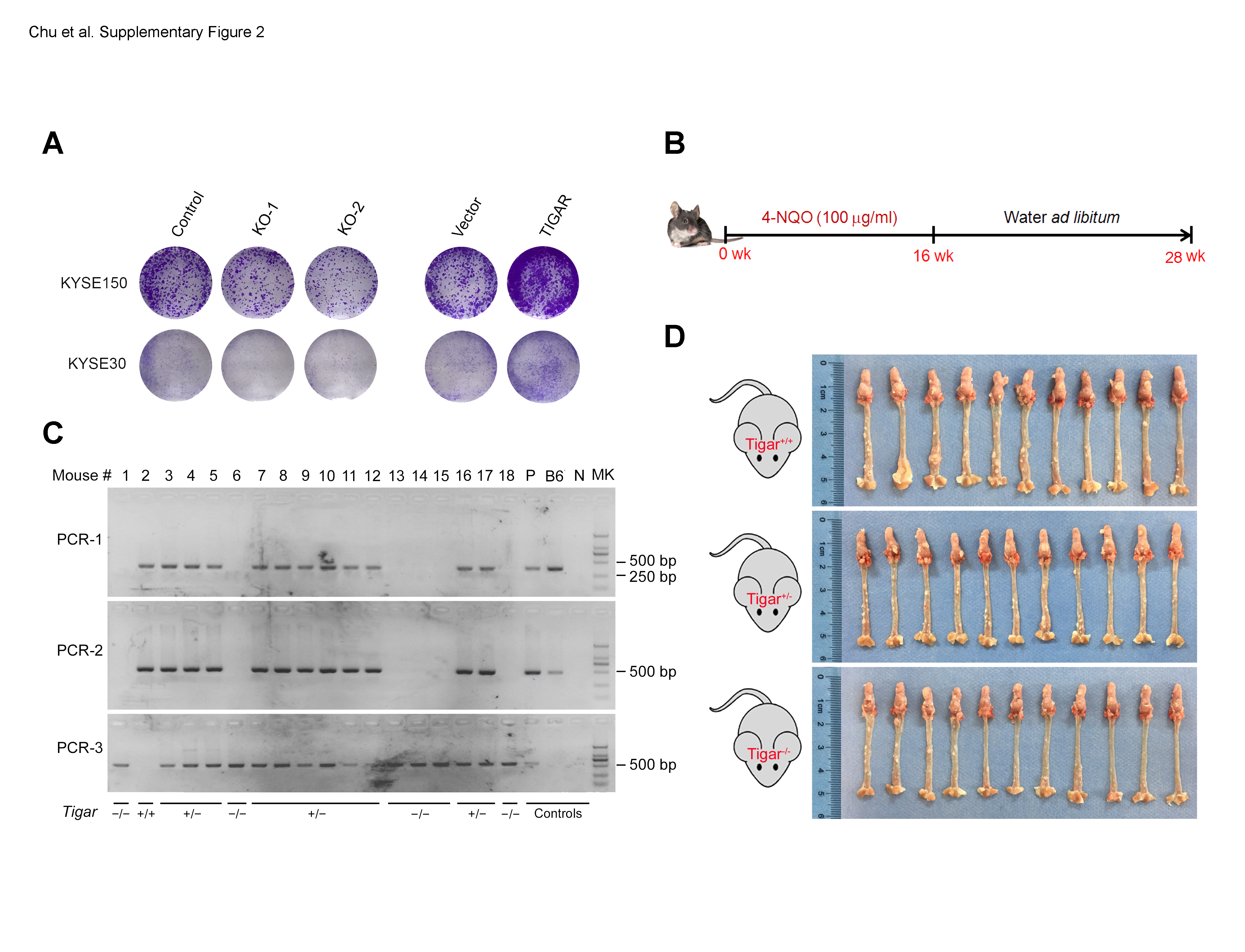

### Supplementary Figue S3

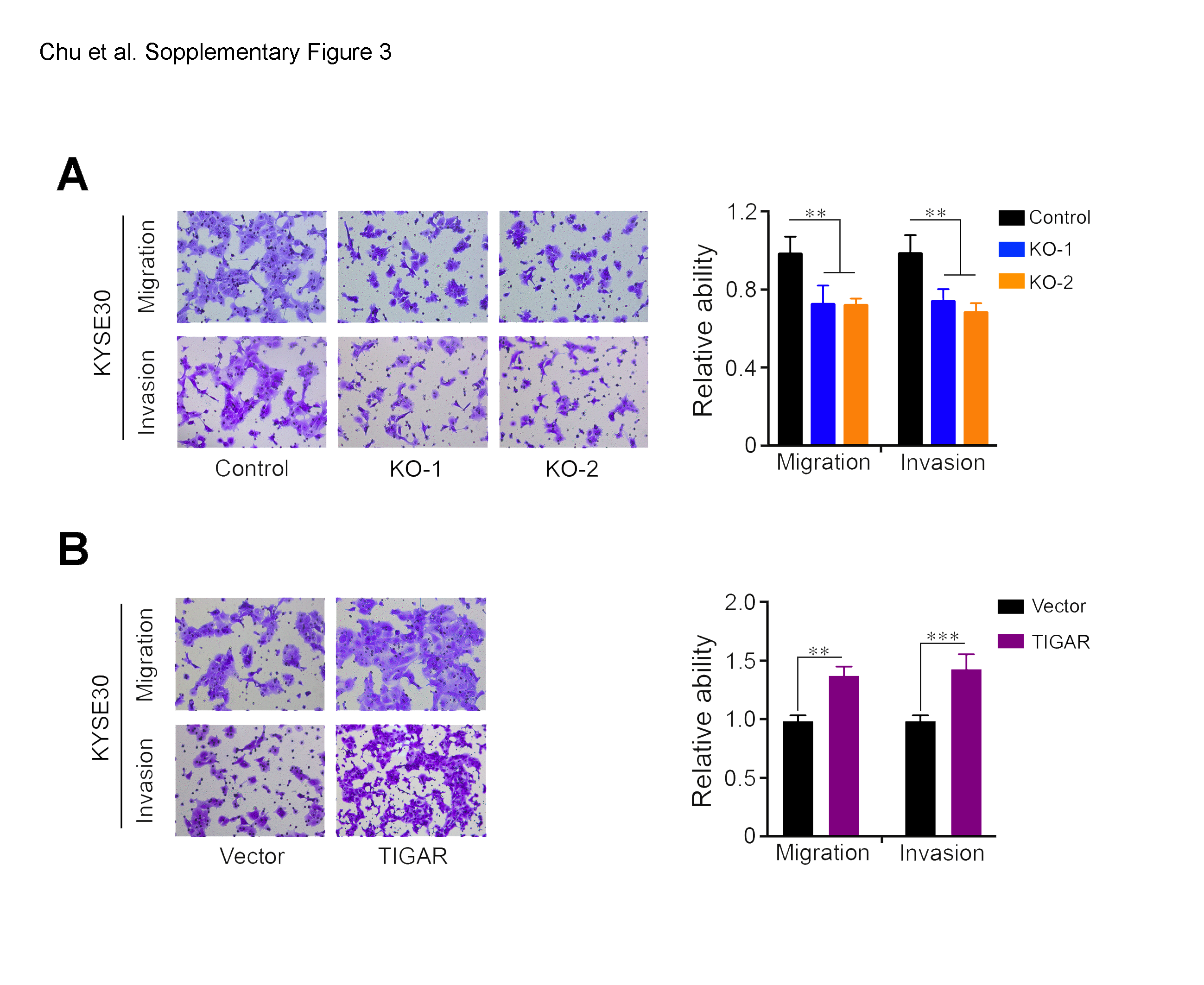

### Supplementary Figue S4

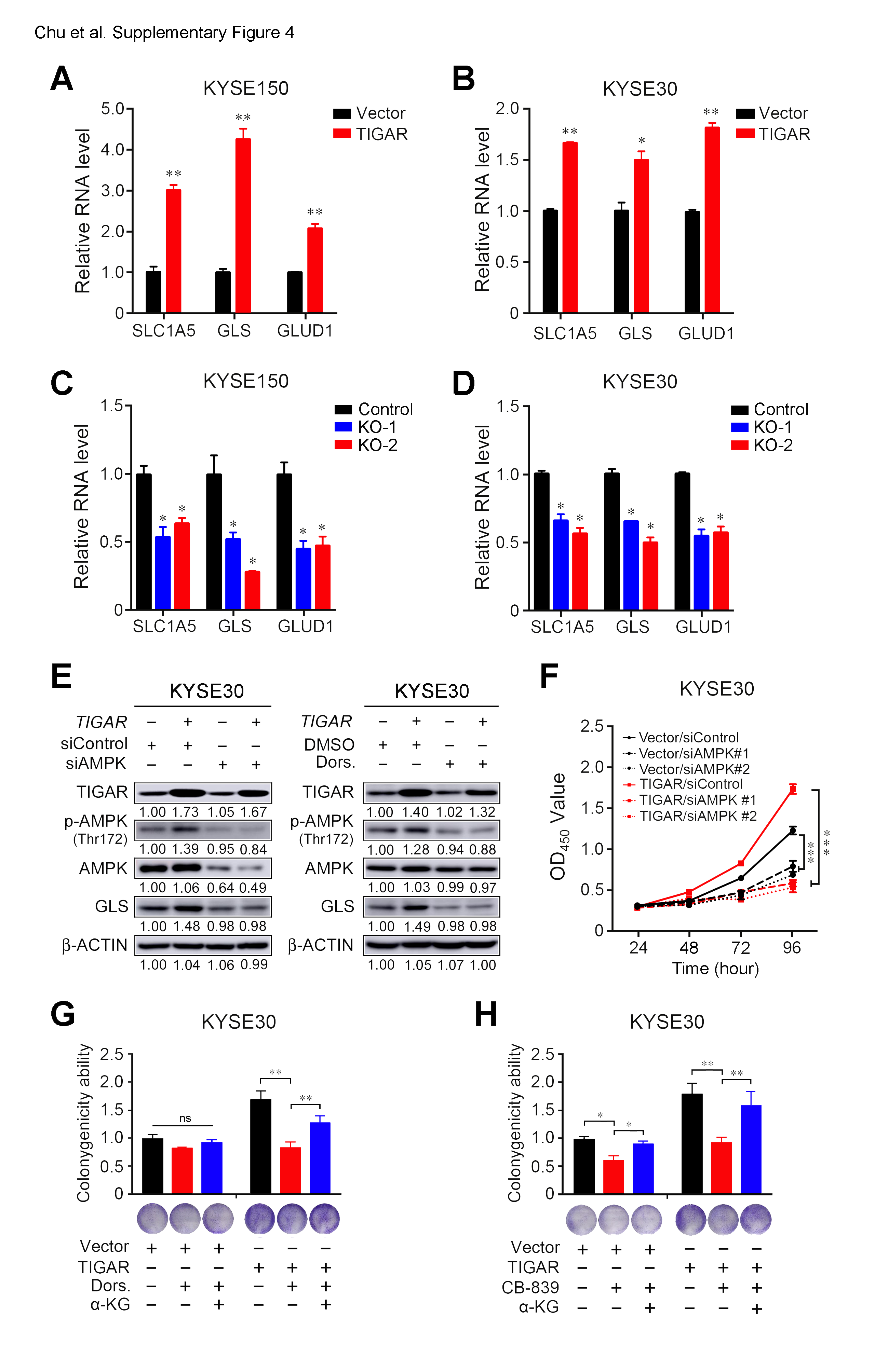

### Supplementary Figue S5

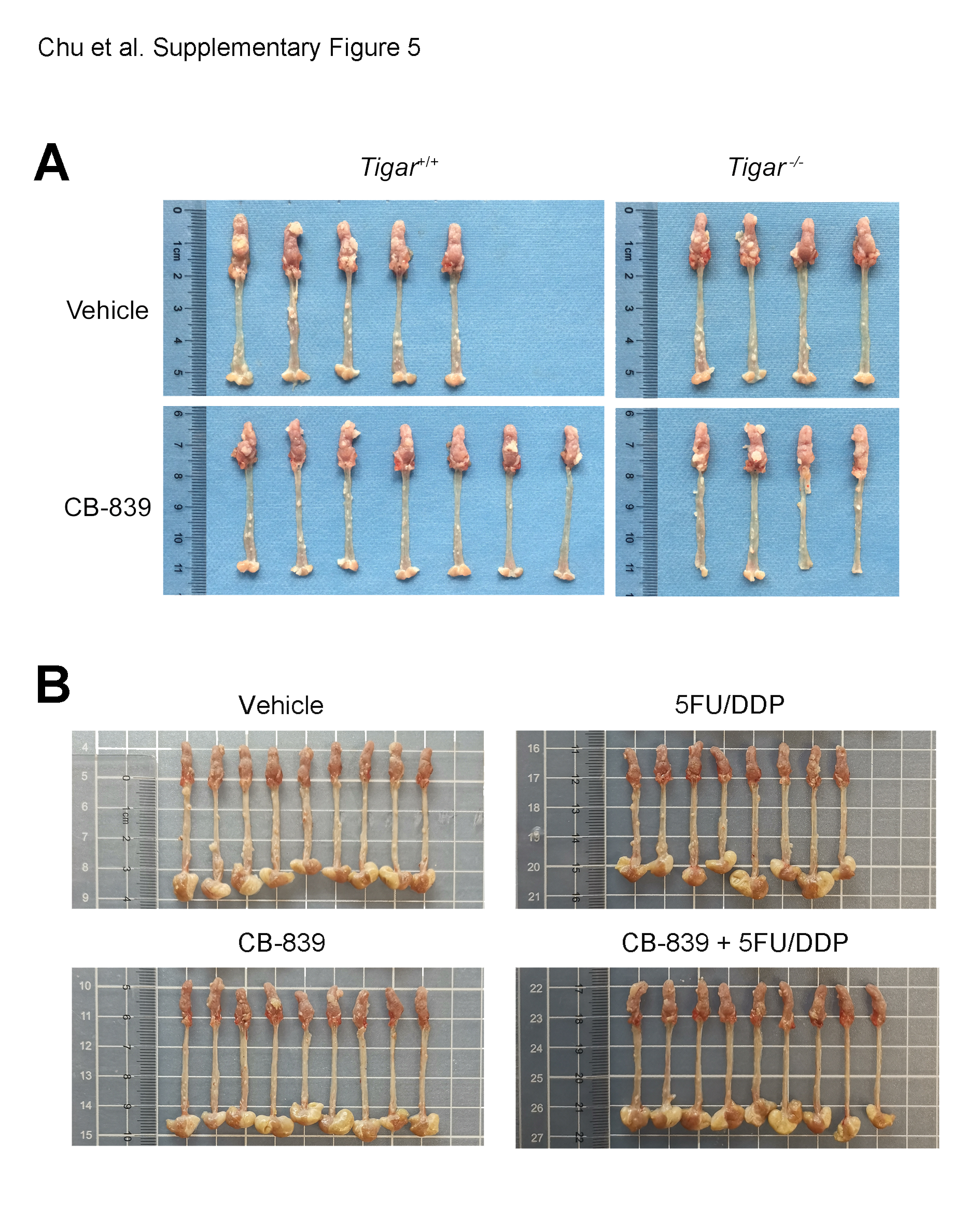

### Supplementary Figue S6

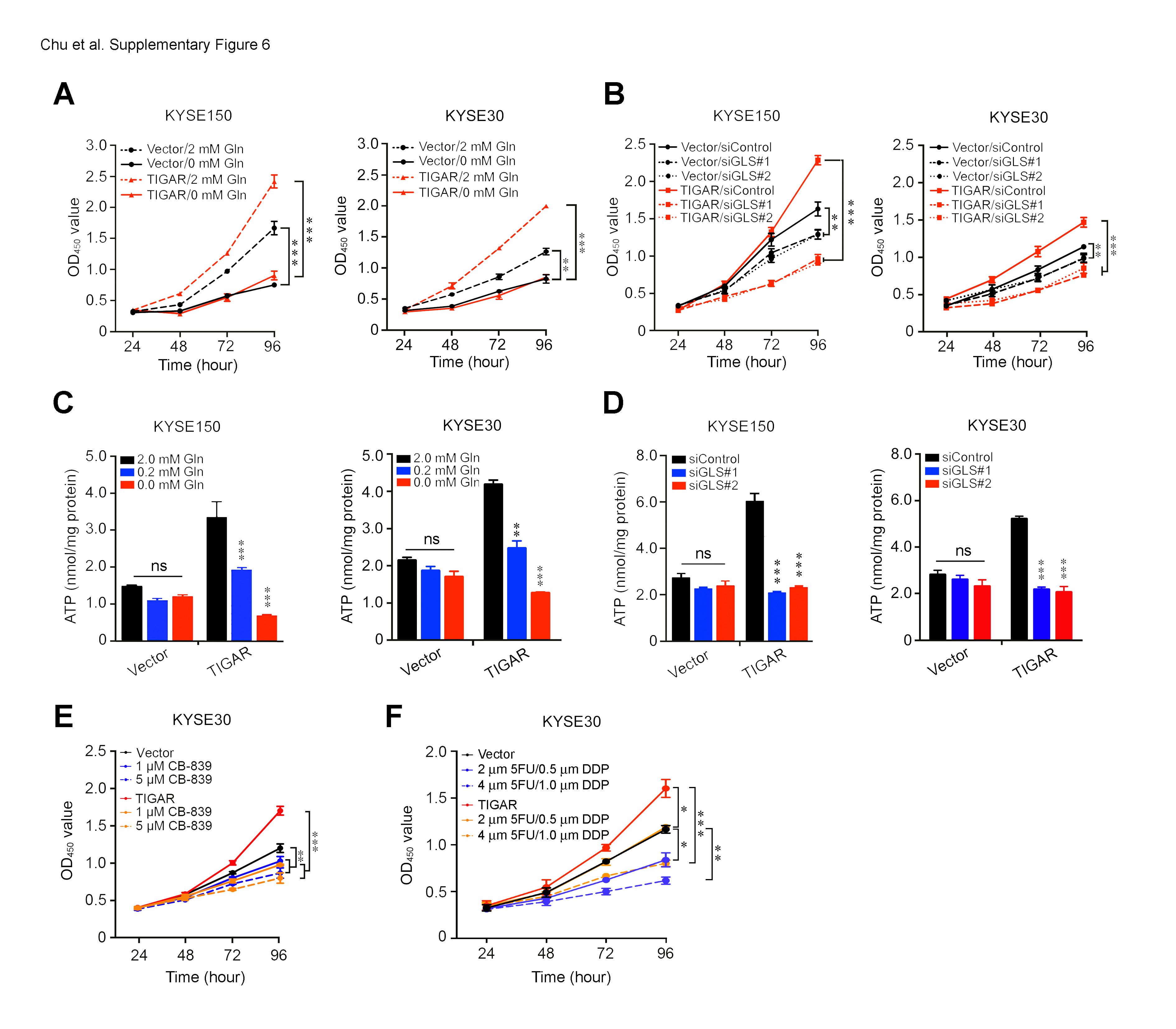

### Supplementary Figue S8

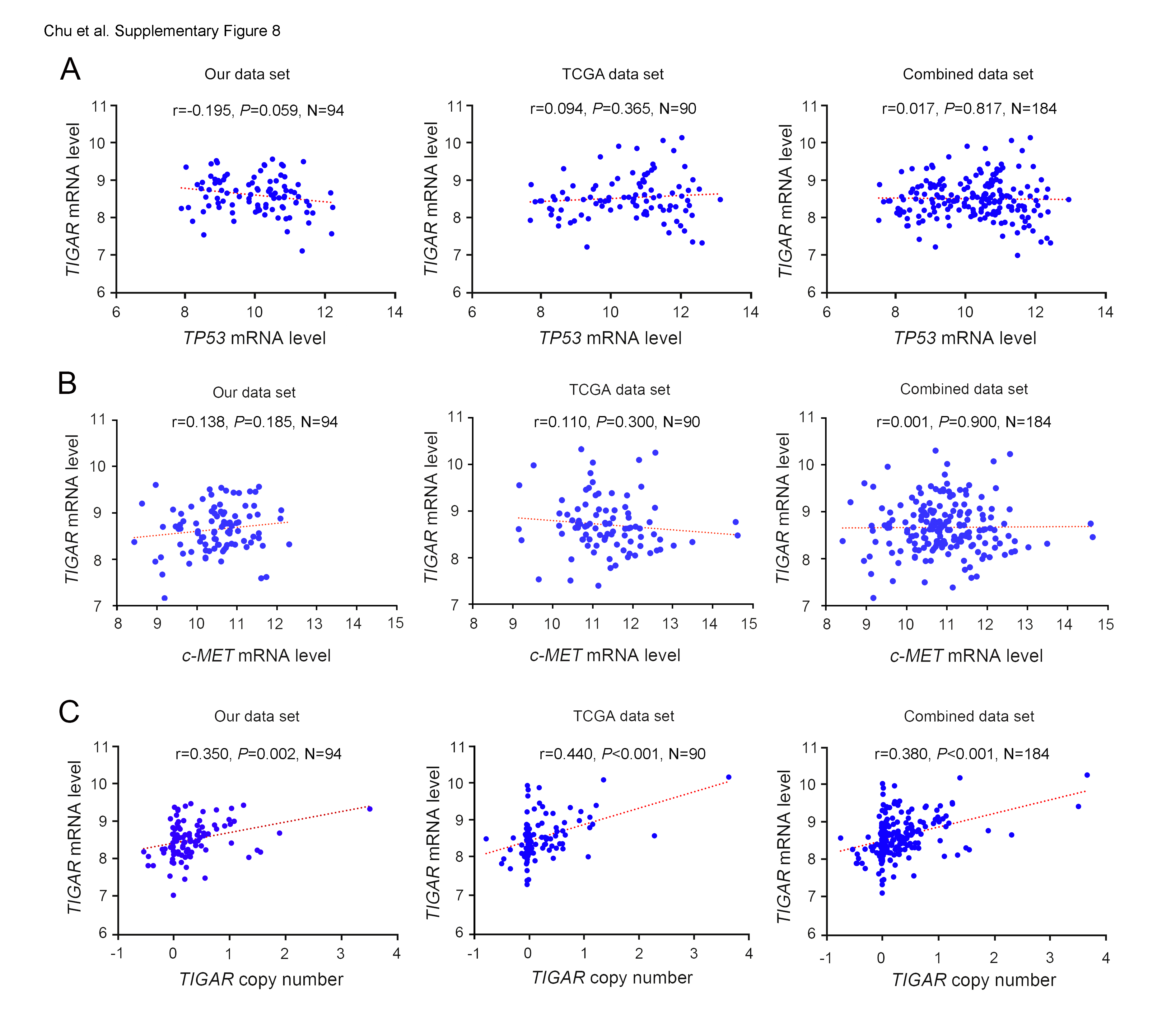

### Supplementary Figue S9

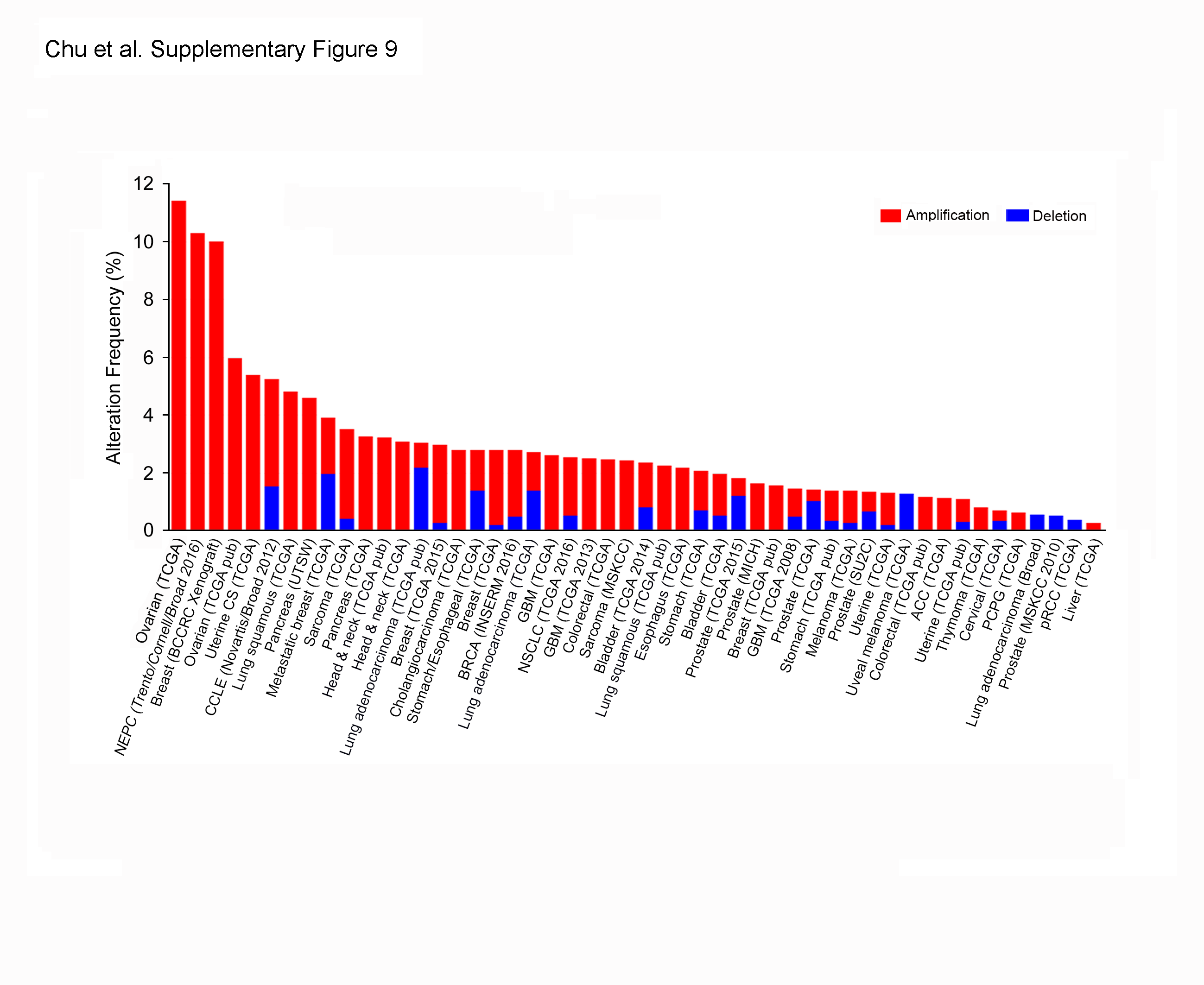
