## Supplementary Figure Legends for "Metabolic remodeling by TIGAR overexpression is a therapeutic target in esophageal squamous-cell carcinoma"

**Supplementary Figure 1. Schematic diagram of data analysis and high-content screening in this study.**

(A) Flow chart for data analysis.

(B) Work flow for high-content screening.

(C) Work flow for the selection of functional genes.

**Supplementary Figure 2. Identification of TIGAR as a player in ESCC proliferation and progression.** (A) Typical pictures of the colony formation of ESCC cells with TIGAR knockout (KO) or overexpression (OE). The quantitative data of colony formation are shown in **Figures 1E** and **2F**.

(B) Schematic diagram of treatment of mice with chemical carcinogen 4-NQO to induce ESCC (see Methods for details).

(C) Genotyping of *Tigar* in *Tigar*-knocked out mice by 3-time PCR. P, positive control for each pair of primers; B6, DNA template from C57BL/6 mice; N, negative control; MK, DNA size markers.

(D) Pictures of the whole esophagus from *Tigar*^+/+^, *Tigar*^+/-^ or *Tigar*^-/-^ mice treated with 4-NQO. The quantitative results of tumor burdens in mice with different genotype are shown in **Figure 1I**.

**Supplementary Figure 3.** **Effects of TIGAR on migration and invasion of ESCC cells in vitro.**

(A) knockout (KO) of *TIGAR* significantly suppressed the migration and invasion abilities of KYSE30 ESCC cells.

(B) Overexpression of *TIGAR* significantly promoted the migration and invasion abilities of KYSE30 ESCC cells.

Left panels are representative pictures of cells in transwell assays and right panels represent data (mean ± SEM) from three independent experiments and each had duplicate. **, *P*<0.01 and ***, *P*<0.001 of Student’s t-test.

**Supplementary Figure 4. TIGAR inhibits glycolysis but activates glutamine pathway via AMPK to promote ESCC.**

(A-D) TIGAR overexpression (A and B) or knockout (KO) (C and D) on the mRNA expression levels of *SLC1A5*, *GLS* and *GLUD1* in ESCC cells. The mRNA levels were determined by qRT-PCR and are represented as mean ± SEM from 3 experiments and each had 3 replications. *, *P*<0.05 and **, *P*<0.01 of two-sided Student’s t-test.

(E) Western blot analysis of GLS in KYSE30 cells with or without TIGAR overexpression and with or without AMPK knockdown (*left*) or inhibition (*right*).

(F) AMPK knockdown significantly repressed KYSE30 cell proliferation and the effect was more pronounced in cells with TIGAR overexpression. Data represent mean ± SEM from 3 experiments and each had 3 replications. ***, *P*<0.001 of two-sided Student’s t-test.

(G and H) Inhibitor of AMPK (Dorsomorphin, Dors.) (G) or GLS (CB-839) (H) significantly repressed KYSE30 cell colony formation and the effect was more pronounced in cells with TIGAR overexpression. α-Ketoglutarate (α-KG), a metabolite of glutamine, significantly rescued ESCC cell colony formation and that was suppressed by AMPK or GLS inhibition. **P*<0.05; **, *P*<0.01; ***, *P*<0.001 of two-sided Student’s t-test of mean ± SEM from 3 experiments and each had 3 replications.

**Supplementary Figure 5.** **Glutaminase inhibitor represses primary ESCC and enhances chemotherapy efficacy in mice.**

(A) Gross anatomy showing ESCC induced by 4-NQO in *Tigar*^+/+^ or *Tigar*^-/-^ mice treated with or without CB-839. Mice were given 4-NQO (see Methods for details) for 16 weeks and after 28weeks, they received CB-839 (200 mg/kg, p. o.; every day) treatment for 18 days.

(B) Gross anatomy showing ESCC induced by 4-NQO in *Tigar*^+/+^ mice treated with CB-839 (200 mg/kg, p.o., every day) or 5FU (20 mg/kg, i. p., every 3 days)/DDP (2 mg/kg, i. p., every 5 days) separately or combination. Mice were given 4-NQO (see Methods for details) for 16 weeks and after 28 weeks, they received the therapies for 21 days.

**Supplementary Figure 6. Inhibition of glutamine pathway inhibits cell proliferation and ATP levels in TIGAR-overexpressed human ESCC cell lines**

(A−D) Glutamine (Gln) deprivation or GLS expression knockdown significantly repressed proliferation (A and B) and ATP production (C and D) in ESCC cells and the effects were more pronounced in cells with TIGAR overexpression. Error bars represent SEM obtained from three independent experiments. **, *P*<0.01 and ***, *P*<0.001 from Student’s t-test.

(E) Dose-dependent inhibition of KYSE30 cell proliferation by CB-839. The effect was more pronounced in cells with TIGAR overexpression. Error bars represent SEM. **, *P*<0.01 and ***, *P*<0.001 from Student’s t-test.

(F) Dose-dependent inhibitory effect of 5FU and DDP treatment on KYSE30 cell proliferation. The efficacy was less pronounced in cells with TIGAR overexpression, suggesting chemoresistance. Error bars represent SEM. ****, *P*<0.0001 from Student’s t-test.

**Supplementary Figure 7.** **Glutamine pathway is a therapeutic target for TIGAR-overexpressed human ESCC.**

(A-D) Hematoxylin and eosin (H&E) staining (*left*) and immunohistochemical staining of TIGAR, p-AMPK, GLS (*middle and right*) in primary tumors of patients and patient-derived xenografts (PDX) in mice with high TIGAR expression (A and B) or low TIGAR expression (C and D).

(E and F) Tumor size in the experiment end from PDX with high TIGAR expression at 19 days (E) and low TIGAR expression at 30 days (F).

(G-J) Combination of 5FU/DDP) and CB-839 treatment significantly repressed mouse PDX with high TIGAR expression as shown in tumor growth curves (G) and tumor size in the experiment end at 21 days (H) and PDX with TIGAR low expression as shown in tumor growth curves (I) and tumor size in the experiment end at 30 days (J). Error bars represent SEM. *, *P*<0.05 from Student’s t-test.

(K) Hematoxylin and eosin (H&E) staining (*top*) and immunohistochemical staining of Ki67 and cleaved CASPASE 3 (*middle* and *bottom*) in PDX with high TIGAR expression, showing that combined 5FU/DDP and CB-839 inhibited proliferation and promoted apoptosis of PDX.

(L) Hematoxylin and eosin (H&E) staining (*top*) and immunohistochemical staining of Ki67 and cleaved CASPASE 3 (*bottom*) in PDX with TIGAR low expression, showing that 5FU/DDP inhibited proliferation and promoted apoptosis of PDX.

**Supplementary Figure 8.** **The *TIGAR* mRNA levels are correlated with *TIGAR* copy number but not *TP53* or *c-MET* mRNA levels in ESCC samples.**

(A) No correlation between *TIGAR* mRNA and *TP53* mRNA levels.

(B) No correlation between *TIGAR* mRNA and *c-MET* mRNA levels.

(C) Positive correlation between *TIGAR* mRNA levels and *TIGAR* gene copy number. Shown are results from our data set, the TCGA data set, and combined data set.

**Supplementary Figure 9.** ***TIGAR* is amplified in many types of human cancer.**

Shown are the frequencies of *TIGAR* alterations in various types of human cancer derived from the cBioportal database, indicating that amplification is a major genomic change.
