## Supplementary Table S1 for "Metabolic remodeling by TIGAR overexpression is a therapeutic target in esophageal squamous-cell carcinoma"

**Supplementary Table S1**. Score of Immunohistochemical Staining of TIGAR, Phosphorylated-AMPK and Glutaminase in Esophageal Squamous-Cell Carcinoma Specimens

| Sample ID | TIGAR | | |  | Phosphorylated-AMPK | | |  | Glutaminase | | |
| --- | --- | --- | --- | --- | --- | --- | --- | --- | --- | --- | --- |
|  | Positive | Intensity | IRS* |  | Positive | Intensity | IRS* |  | Positive | Intensity | IRS* |
| 160408 | 2 | 2 | 4 |  | 2 | 1 | 2 |  | 2 | 2 | 4 |
| 160425 | 2 | 2 | 4 |  | 2 | 2 | 4 |  | 2 | 1 | 2 |
| 160442 | 2 | 3 | 6 |  | 2 | 2 | 4 |  | 2 | 3 | 6 |
| 160456 | 3 | 2 | 6 |  | 2 | 2 | 4 |  | 2 | 3 | 6 |
| 160466 | 3 | 3 | 9 |  | 2 | 3 | 6 |  | 4 | 3 | 12 |
| 160474 | 2 | 2 | 4 |  | 2 | 2 | 4 |  | 3 | 3 | 9 |
| 160475 | 4 | 2 | 8 |  | 3 | 3 | 9 |  | 3 | 2 | 6 |
| 160467 | 3 | 3 | 9 |  | 2 | 2 | 4 |  | 3 | 3 | 9 |
| 160458 | 3 | 3 | 9 |  | 3 | 3 | 9 |  | 4 | 3 | 12 |
| 160443 | 3 | 2 | 6 |  | 2 | 2 | 4 |  | 3 | 3 | 9 |
| 160430 | 2 | 3 | 6 |  | 2 | 1 | 2 |  | 2 | 2 | 4 |
| 160413 | 0 | 0 | 0 |  | 2 | 1 | 2 |  | 2 | 1 | 2 |
| 160432 | 3 | 2 | 6 |  | 2 | 2 | 4 |  | 2 | 2 | 4 |
| 160460 | 2 | 2 | 4 |  | 2 | 2 | 4 |  | 2 | 3 | 6 |
| 160468 | 2 | 1 | 2 |  | 2 | 2 | 4 |  | 2 | 3 | 6 |
| 160469 | 2 | 3 | 6 |  | 2 | 2 | 4 |  | 2 | 2 | 4 |
| 160461 | 3 | 1 | 3 |  | 2 | 1 | 2 |  | 2 | 1 | 2 |
| 160453 | 2 | 3 | 6 |  | 2 | 2 | 4 |  | 3 | 3 | 9 |
| 160414 | 2 | 1 | 2 |  | 2 | 1 | 2 |  | 2 | 2 | 4 |
| 160421 | 2 | 2 | 4 |  | 2 | 1 | 2 |  | 2 | 2 | 4 |
| 160439 | 2 | 3 | 6 |  | 2 | 3 | 6 |  | 2 | 3 | 6 |
| 160454 | 2 | 2 | 4 |  | 2 | 1 | 2 |  | 2 | 2 | 4 |
| 160470 | 3 | 3 | 9 |  | 2 | 2 | 4 |  | 2 | 3 | 6 |
| 160473 | 2 | 2 | 4 |  | 2 | 3 | 6 |  | 2 | 3 | 6 |
| 160463 | 3 | 1 | 3 |  | 2 | 3 | 6 |  | 4 | 3 | 12 |
| 160455 | 2 | 2 | 4 |  | 2 | 2 | 4 |  | 2 | 2 | 4 |
| 160441 | 2 | 3 | 6 |  | 2 | 3 | 6 |  | 3 | 3 | 9 |
| 160422 | 4 | 3 | 12 |  | 3 | 3 | 9 |  | 4 | 3 | 12 |

*IRS, immune reactive score. IRS was obtained by multiplying the score of positive and that of intensity. The labeling score of positive was defined as 1 (≤10%), 2 (11%−50%), 3 (51%−80%) and 4 (>80%); the labeling score of intensity was estimated as negative (0), weak (1), moderate (2) and strong (3).
