## Supplementary Table S2 for "Metabolic remodeling by TIGAR overexpression is a therapeutic target in esophageal squamous-cell carcinoma"

**Supplementary Table S2**. Characteristics of 225 Patients with Esophageal Squamous-Cell Carcinoma in This Study

| Sample ID | Age  (year) | Sex* | Smoker^#^ | Drinker^#^ | TNM^§^ stage | T stage | N stage | M stage | Survival status | Survival time (month) |
| --- | --- | --- | --- | --- | --- | --- | --- | --- | --- | --- |
| 150113 | 70 | M | Yes | No | III | 3 | 1 | 0 | Deceased | Missing |
| 150116 | 72 | M | Yes | No | III | 3 | 2 | 0 | Deceased | 12.4 |
| 150133 | 64 | M | Yes | Yes | II | 2 | 0 | 0 | Alive | 46.2 |
| 150146 | 52 | M | Yes | Yes | II | 3 | 0 | 0 | Deceased | 25.1 |
| 150152 | 61 | M | Yes | Yes | III | 3 | 3 | 0 | Deceased | Missing |
| 150159 | 63 | M | Yes | No | II | 2 | 0 | 0 | Alive | 45.8 |
| 150201 | 63 | M | No | No | II | 3 | 0 | 0 | Alive | 45.7 |
| 150203 | 61 | M | Yes | No | I | 3 | 0 | 0 | Deceased | 38.6 |
| 150208 | 60 | M | Yes | Yes | III | 2 | 2 | 0 | Deceased | 17.6 |
| 150219 | 68 | M | No | No | III | 3 | 3 | 0 | Deceased | 21.6 |
| 150228 | 61 | M | Yes | Yes | I | 1 | 0 | 0 | Alive | 45.4 |
| 150229 | 67 | F | No | No | II | 2 | 0 | 0 | Alive | 45.3 |
| 150233 | 53 | M | Yes | Yes | III | 3 | 1 | 0 | Deceased | 11.7 |
| 150234 | 61 | M | Yes | No | II | 2 | 0 | 0 | Alive | 44.8 |
| 150241 | 62 | M | Yes | Yes | III | 3 | 2 | 0 | Deceased | 10.0 |
| 150302 | 62 | M | No | No | III | 3 | 1 | 0 | Deceased | 20.8 |
| 150304 | 58 | M | Yes | No | II | 2 | 1 | 0 | Deceased | 23.9 |
| 150311 | 63 | M | Yes | Yes | III | 3 | 2 | 0 | Deceased | Missing |
| 150312 | 62 | M | Yes | Yes | II | 3 | 0 | 0 | Alive | 44.9 |
| 150313 | 69 | M | Yes | No | II | 3 | 0 | 0 | Alive | 44.9 |
| 150319 | 77 | M | Yes | No | II | 2 | 1 | 0 | Deceased | 3.4 |
| 150322 | 70 | M | No | No | II | 3 | 0 | 0 | Deceased | 32.9 |
| 150323 | 66 | M | No | No | III | 3 | 1 | 0 | Deceased | 31.9 |
| 150324 | 51 | M | Yes | Yes | II | 2 | 0 | 0 | Alive | 44.8 |
| 150326 | 60 | M | Yes | Yes | II | 3 | 0 | 0 | Deceased | Missing |
| 150331 | 69 | M | Yes | Yes | II | 2 | 1 | 0 | Deceased | 19.6 |
| 150333 | 69 | M | Yes | No | II | 3 | 0 | 0 | Deceased | 27.6 |
| 150334 | 56 | M | Yes | No | IV | 3 | 2 | 1 | Deceased | 30.8 |
| 150339 | 55 | M | Yes | Yes | II | 2 | 0 | 0 | Alive | 44.7 |
| 150347 | 73 | M | No | No | II | 2 | 0 | 0 | Deceased | Missing |
| 150348 | 74 | M | Yes | Yes | I | 1 | 0 | 0 | Deceased | 33.7 |
| 150355 | 58 | M | Yes | Yes | III | 3 | 1 | 0 | Deceased | 0.5 |
| 150374 | 57 | F | No | No | II | 2 | 1 | 0 | Alive | 44.4 |
| 150378 | 60 | M | No | No | I | 1 | 0 | 0 | Alive | 44.4 |
| 150385 | 71 | F | No | No | III | 3 | 2 | 0 | Alive | 44.3 |
| 150388 | 65 | M | Yes | Yes | II | 3 | 0 | 0 | Deceased | 7.9 |
| 150389 | 64 | M | Yes | Yes | I | 1 | 0 | 0 | Deceased | Missing |
| 150395 | 62 | F | No | No | III | 3 | 1 | 0 | Deceased | 11.9 |
| 150411 | 50 | M | Yes | Yes | II | 3 | 0 | 0 | Alive | 43.8 |
| 150414 | 65 | M | Yes | Yes | II | 2 | 0 | 0 | Alive | 43.8 |
| 150418 | 65 | M | Yes | Yes | III | 3 | 2 | 0 | Deceased | 6.6 |
| 150419 | 58 | F | No | No | III | 3 | 1 | 0 | Alive | 43.7 |
| 150423 | 62 | M | Yes | Yes | II | 3 | 0 | 0 | Alive | 43.7 |
| 150429 | 52 | M | Yes | Yes | III | 3 | 3 | 0 | Deceased | 7.3 |
| 150431 | 59 | M | Yes | Yes | II | 2 | 0 | 0 | Alive | 43.6 |
| 150432 | 57 | F | No | No | II | 3 | 0 | 0 | Alive | 43.5 |
| 150434 | 62 | M | Yes | Yes | III | 3 | 3 | 0 | Deceased | Missing |
| 150436 | 76 | M | Yes | Yes | IV | 3 | 2 | 1 | Deceased | 24.4 |
| 150441 | 50 | M | Yes | Yes | II | 2 | 1 | 0 | Alive | 42.5 |
| 150454 | 62 | M | No | No | III | 3 | 2 | 0 | Alive | 42.3 |
| 150455 | 65 | M | No | Yes | III | 4 | 1 | 0 | Deceased | 4.9 |
| 150458 | 68 | F | No | No | II | 3 | 0 | 0 | Deceased | 34.3 |
| 150459 | 59 | F | No | No | II | 3 | 0 | 0 | Alive | 42.2 |
| 150468 | 64 | M | Yes | Yes | IV | 3 | 2 | 1 | Deceased | 32.1 |
| 150472 | 74 | M | No | No | III | 3 | 1 | 0 | Deceased | 27.0 |
| 150473 | 58 | M | No | Yes | I | 3 | 0 | 0 | Alive | 42.1 |
| 150474 | 68 | M | Yes | No | III | 3 | 1 | 0 | Alive | 42.1 |
| 150476 | 60 | M | No | No | II | 2 | 0 | 0 | Alive | 42.0 |
| 150481 | 65 | F | No | No | IV | 3 | 2 | 1 | Alive | 41.8 |
| 150482 | 70 | M | Yes | Yes | II | 3 | 0 | 0 | Deceased | 2.5 |
| 150502 | 52 | M | Yes | Yes | III | 3 | 1 | 0 | Deceased | 7.5 |
| 150506 | 73 | M | Yes | No | IV | 3 | 2 | 1 | Deceased | 4.4 |
| 150510 | 55 | M | Yes | Yes | II | 3 | 0 | 0 | Alive | 41.8 |
| 150513 | 61 | M | Yes | No | II | 3 | 0 | 0 | Deceased | 3.5 |
| 150526 | 60 | M | Yes | Yes | II | 2 | 0 | 0 | Alive | 41.6 |
| 150527 | 67 | F | No | No | II | 2 | 1 | 0 | Deceased | 7.2 |
| 150528 | 68 | M | Yes | Yes | II | 3 | 0 | 0 | Alive | 41.6 |
| 150531 | 57 | M | Yes | Yes | III | 2 | 2 | 0 | Alive | 41.8 |
| 150541 | 69 | M | Yes | Yes | I | 1 | 0 | 0 | Alive | 41.7 |
| 150543 | 74 | M | Yes | Yes | III | 3 | 1 | 0 | Deceased | 14.1 |
| 150555 | 66 | M | Yes | No | II | 3 | 0 | 0 | Deceased | 23.7 |
| 150557 | 64 | F | No | No | II | 2 | 0 | 0 | Alive | 41.5 |
| 150560 | 70 | M | Yes | Yes | II | 3 | 0 | 0 | Deceased | 32.9 |
| 150561 | 58 | M | Yes | No | III | 3 | 1 | 0 | Deceased | 9.0 |
| 150582 | 48 | F | No | No | I | 1 | 0 | 0 | Alive | 41.2 |
| 150585 | 65 | M | Yes | Yes | III | 2 | 2 | 0 | Deceased | 1.6 |
| 150601 | 64 | M | Yes | No | III | 4 | 0 | 0 | Alive | 41.1 |
| 150603 | 78 | M | Yes | Yes | III | 4 | 1 | 0 | Alive | 41.1 |
| 150606 | 58 | F | No | No | III | 3 | 1 | 0 | Deceased | 10.6 |
| 150609 | 49 | M | Yes | Yes | III | 3 | 1 | 0 | Deceased | 12.6 |
| 150613 | 53 | F | No | No | II | 1 | 1 | 0 | Alive | 41.0 |
| 150614 | 67 | M | Yes | Yes | III | 3 | 1 | 0 | Deceased | 1.9 |
| 150617 | 74 | F | No | No | II | 2 | 0 | 0 | Deceased | 28.8 |
| 150618 | 52 | M | Yes | Yes | II | 3 | 0 | 0 | Alive | 40.9 |
| 150622 | 75 | M | No | No | II | 2 | 0 | 0 | Deceased | 21.5 |
| 150623 | 62 | M | No | Yes | II | 3 | 0 | 0 | Deceased | 6.3 |
| 150627 | 58 | M | Yes | No | III | 3 | 3 | 0 | Deceased | 26.6 |
| 150631 | 61 | M | Yes | Yes | II | 3 | 0 | 0 | Alive | 40.8 |
| 150641 | 52 | M | Yes | Yes | III | 3 | 1 | 0 | Deceased | 6.0 |
| 150645 | 71 | F | No | No | II | 2 | 0 | 0 | Deceased | 12.6 |
| 150654 | 61 | M | Yes | Yes | III | 3 | 1 | 0 | Alive | 40.5 |
| 150660 | 71 | F | No | No | III | 2 | 2 | 0 | Deceased | 15.7 |
| 150662 | 57 | M | No | No | II | 1 | 1 | 0 | Alive | 40.4 |
| 150664 | 58 | M | Yes | Yes | III | 2 | 2 | 0 | Deceased | 12.2 |
| 150666 | 48 | M | Yes | Yes | I | 1 | 0 | 0 | Deceased | 6.2 |
| 150667 | 56 | M | Yes | Yes | II | 1 | 1 | 0 | Deceased | 22.8 |
| 150671 | 65 | F | No | No | II | 2 | 0 | 0 | Alive | 40.2 |
| 150703 | 44 | M | No | No | II | 2 | 1 | 0 | Alive | 40.1 |
| 150705 | 63 | M | Yes | No | I | 2 | 0 | 0 | Alive | 40.1 |
| 150707 | 59 | M | Yes | Yes | II | 3 | 0 | 0 | Alive | 40.1 |
| 150710 | 67 | F | No | No | II | 2 | 0 | 0 | Alive | 40.0 |
| 150724 | 71 | M | Yes | No | II | 3 | 0 | 0 | Alive | 39.9 |
| 150726 | 69 | M | Yes | Yes | II | 2 | 0 | 0 | Deceased | 9.3 |
| 150730 | 65 | M | Yes | No | I | 1 | 0 | 0 | Alive | 39.8 |
| 150734 | 75 | F | No | No | II | 2 | 1 | 0 | Alive | 39.7 |
| 150735 | 59 | M | Yes | No | III | 2 | 2 | 0 | Deceased | 22.8 |
| 150736 | 60 | M | Yes | No | III | 3 | 1 | 0 | Alive | 39.7 |
| 150739 | 70 | M | Yes | Yes | II | 2 | 0 | 0 | Deceased | 7.9 |
| 150741 | 73 | M | Yes | No | III | 4 | 3 | 0 | Deceased | 21.4 |
| 150745 | 64 | F | No | No | III | 2 | 3 | 0 | Alive | 39.5 |
| 150747 | 64 | M | Yes | Yes | II | 3 | 0 | 0 | Alive | 39.5 |
| 150749 | 71 | M | Yes | No | III | 3 | 2 | 0 | Alive | 39.5 |
| 150758 | 53 | M | Yes | No | III | 3 | 2 | 0 | Deceased | 2.7 |
| 150762 | 59 | M | Yes | Yes | III | 3 | 1 | 0 | Deceased | 37.6 |
| 150770 | 73 | F | No | No | II | 3 | 0 | 0 | Deceased | 10.5 |
| 150801 | 63 | M | Yes | Yes | III | 3 | 1 | 0 | Deceased | 2.4 |
| 150805 | 51 | F | No | No | II | 3 | 0 | 0 | Alive | 39.0 |
| 150806 | 61 | M | Yes | Yes | II | 3 | 0 | 0 | Alive | 39.0 |
| 150807 | 64 | M | No | Yes | III | 1 | 3 | 0 | Deceased | 9.0 |
| 150812 | 52 | M | Yes | Yes | III | 3 | 1 | 0 | Deceased | 5.4 |
| 150813 | 50 | M | No | No | I | 2 | 0 | 0 | Deceased | 1.6 |
| 150843 | 58 | M | Yes | Yes | III | 3 | 1 | 0 | Deceased | 23.0 |
| 150845 | 66 | F | No | No | I | 1 | 0 | 0 | Alive | 38.3 |
| 150846 | 74 | M | No | No | III | 3 | 1 | 0 | Alive | 38.3 |
| 150848 | 53 | F | No | No | III | 3 | 1 | 0 | Alive | 38.3 |
| 150854 | 60 | F | No | No | II | 3 | 0 | 0 | Deceased | 28.0 |
| 150901 | 57 | F | No | No | I | 1 | 0 | 0 | Alive | 38.0 |
| 150903 | 51 | M | Yes | Yes | III | 3 | 1 | 0 | Alive | 38.1 |
| 150906 | 55 | M | Yes | No | III | 3 | 2 | 0 | Alive | 37.9 |
| 150909 | 72 | M | Yes | No | II | 2 | 0 | 0 | Alive | 37.9 |
| 150914 | 63 | M | Yes | Yes | III | 3 | 1 | 0 | Deceased | 5.3 |
| 150917 | 72 | M | Yes | No | II | 3 | 0 | 0 | Deceased | Missing |
| 150918 | 63 | F | No | No | III | 2 | 3 | 0 | Alive | 37.6 |
| 150923 | 74 | M | Yes | No | II | 3 | 0 | 0 | Deceased | 23.5 |
| 150933 | 51 | M | Yes | Yes | II | 3 | 0 | 0 | Alive | 37.4 |
| 150934 | 61 | M | Yes | Yes | I | 3 | 0 | 0 | Alive | 37.4 |
| 150935 | 61 | F | No | No | III | 3 | 1 | 0 | Alive | 37.4 |
| 150936 | 65 | M | Yes | Yes | II | 3 | 0 | 0 | Alive | 37.4 |
| 150947 | 61 | M | No | Yes | II | 3 | 0 | 0 | Alive | 37.5 |
| 150953 | 68 | M | Yes | Yes | III | 3 | 1 | 0 | Deceased | 5.8 |
| 150954 | 65 | F | No | No | II | 3 | 0 | 0 | Alive | 37.5 |
| 151002 | 58 | F | No | No | II | 3 | 0 | 0 | Alive | 37.2 |
| 151011 | 67 | M | Yes | No | III | 3 | 1 | 0 | Deceased | 4.3 |
| 151012 | 69 | F | No | No | II | 3 | 0 | 0 | Deceased | 32.7 |
| 151016 | 63 | M | Yes | Yes | II | 2 | 0 | 0 | Deceased | 34.3 |
| 151020 | 68 | F | No | No | II | 2 | 0 | 0 | Alive | 37.0 |
| 151021 | 62 | M | Yes | Yes | II | 3 | 0 | 0 | Deceased | 15.8 |
| 151024 | 55 | F | No | No | II | 3 | 0 | 0 | Alive | 36.9 |
| 151033 | 50 | M | Yes | Yes | II | 3 | 0 | 0 | Alive | 36.7 |
| 151040 | 67 | M | No | No | II | 2 | 1 | 0 | Alive | 36.7 |
| 151044 | 52 | M | Yes | Yes | III | 3 | 1 | 0 | Deceased | 25.2 |
| 151047 | 65 | M | Yes | Yes | II | 3 | 0 | 0 | Alive | 36.5 |
| 151049 | 52 | M | Yes | Yes | III | 3 | 1 | 0 | Alive | 36.5 |
| 151050 | 46 | M | Yes | Yes | II | 3 | 0 | 0 | Alive | 36.5 |
| 151052 | 61 | M | Yes | Yes | II | 2 | 1 | 0 | Deceased | 19.9 |
| 151053 | 56 | F | No | No | I | 1 | 0 | 0 | Alive | 36.5 |
| 151107 | 68 | M | No | No | III | 3 | 3 | 0 | Alive | 36.5 |
| 151116 | 74 | M | Yes | Yes | III | 3 | 1 | 0 | Alive | 36.4 |
| 151131 | 69 | F | No | Yes | III | 3 | 1 | 0 | Deceased | 0.6 |
| 151133 | 67 | F | No | No | III | 3 | 1 | 0 | Alive | 36.0 |
| 151138 | 49 | M | No | No | II | 3 | 0 | 0 | Alive | 36.0 |
| 151202 | 60 | F | No | No | III | 3 | 1 | 0 | Deceased | Missing |
| 151209 | 52 | M | No | No | II | 2 | 1 | 0 | Deceased | 8.4 |
| 151212 | 70 | M | No | No | II | 2 | 1 | 0 | Deceased | 14.5 |
| 151243 | 63 | F | No | No | II | 3 | 0 | 0 | Alive | 34.1 |
| 151256 | 60 | M | Yes | Yes | III | 3 | 1 | 0 | Deceased | 0.1 |
| 160103 | 61 | M | Yes | Yes | I | 1 | 0 | 0 | Alive | 33.7 |
| 160105 | 66 | F | No | No | II | 2 | 0 | 0 | Alive | 33.7 |
| 160106 | 56 | F | No | No | III | 3 | 1 | 0 | Alive | 33.7 |
| 160110 | 62 | F | No | No | I | 3 | 0 | 0 | Alive | 33.8 |
| 160111 | 66 | F | No | Yes | II | 3 | 0 | 0 | Deceased | 20.6 |
| 160114 | 72 | F | No | No | II | 3 | 0 | 0 | Alive | 33.8 |
| 160117 | 75 | M | Yes | Yes | III | 3 | 1 | 0 | Deceased | 14.4 |
| 160119 | 67 | M | No | No | I | 1 | 0 | 0 | Alive | 33.8 |
| 160131 | 57 | M | Yes | No | III | 3 | 2 | 0 | Deceased | 19.8 |
| 160139 | 65 | M | Yes | Yes | I | 1 | 0 | 0 | Alive | 33.5 |
| 160140 | 56 | M | Yes | Yes | III | 3 | 2 | 0 | Deceased | 23.4 |
| 160143 | 62 | M | Yes | Yes | II | 2 | 0 | 0 | Alive | 33.4 |
| 160151 | 53 | M | Yes | Yes | II | 3 | 0 | 0 | Alive | 33.3 |
| 160161 | 59 | M | Yes | No | IV | 3 | 2 | 1 | Deceased | 17.8 |
| 160204 | 64 | M | Yes | No | II | 2 | 1 | 0 | Alive | 32.6 |
| 160205 | 62 | M | Yes | Yes | I | 1 | 0 | 0 | Alive | 32.3 |
| 160209 | 57 | M | Yes | No | II | 2 | 1 | 0 | Alive | 32.6 |
| 160211 | 72 | M | Yes | Yes | II | 3 | 0 | 0 | Alive | 32.6 |
| 160218 | 64 | M | Yes | No | I | 1 | 0 | 0 | Alive | 32.5 |
| 160219 | 66 | M | Yes | No | III | 3 | 3 | 0 | Deceased | 20.1 |
| 160220 | 61 | F | No | No | III | 4 | 2 | 0 | Deceased | 24.9 |
| 160222 | 52 | M | Yes | Yes | II | 3 | 0 | 0 | Alive | 32.5 |
| 160227 | 72 | F | No | No | I | 1 | 0 | 0 | Alive | 32.5 |
| 160228 | 71 | M | Yes | No | III | 3 | 1 | 0 | Alive | 32.5 |
| 160236 | 62 | M | Yes | Yes | III | 3 | 3 | 0 | Deceased | 18.9 |
| 160238 | 63 | M | No | Yes | III | 3 | 2 | 0 | Deceased | 12.8 |
| 160246 | 70 | F | No | No | II | 3 | 0 | 0 | Alive | 32.3 |
| 160248 | 49 | M | Yes | Yes | II | 3 | 0 | 0 | Alive | 32.3 |
| 160301 | 67 | F | No | No | III | 3 | 1 | 0 | Deceased | 7.6 |
| 160303 | 55 | M | Yes | Yes | III | 3 | 1 | 0 | Alive | 32.3 |
| 160309 | 63 | M | Yes | No | II | 3 | 0 | 0 | Alive | 32.2 |
| 160310 | 69 | M | Yes | Yes | II | 3 | 0 | 0 | Deceased | 9.5 |
| 160311 | 52 | M | Yes | Yes | III | 3 | 1 | 0 | Deceased | 18.8 |
| 160319 | 57 | M | Yes | Yes | III | 3 | 2 | 0 | Deceased | 21.6 |
| 160320 | 64 | M | Yes | Yes | II | 3 | 0 | 0 | Alive | 32.2 |
| 160324 | 53 | M | Yes | Yes | II | 3 | 0 | 0 | Deceased | 16.4 |
| 160326 | 65 | M | Yes | Yes | II | 3 | 0 | 0 | Alive | 32.0 |
| 160333 | 61 | M | Yes | Yes | III | 2 | 3 | 0 | Deceased | 13.3 |
| 160334 | 64 | M | Yes | No | II | 3 | 0 | 0 | Alive | 31.9 |
| 160343 | 66 | M | Yes | Yes | II | 3 | 0 | 0 | Alive | 31.8 |
| 160372 | 53 | F | No | No | I | 1 | 0 | 0 | Alive | 31.4 |
| 160375 | 60 | F | No | No | III | 3 | 1 | 0 | Alive | 31.3 |
| 160413 | 57 | M | No | Yes | III | 3 | 1 | 0 | Deceased | 19.6 |
| 160414 | 62 | M | No | No | I | 1 | 0 | 0 | Alive | 31.2 |
| 160421 | 65 | F | No | No | I | 1 | 0 | 0 | Alive | 31.1 |
| 160430 | 71 | M | No | No | III | 3 | 2 | 0 | Deceased | 18.5 |
| 160432 | 73 | M | Yes | No | III | 3 | 3 | 0 | Deceased | 5.6 |
| 160439 | 66 | M | Yes | Yes | II | 3 | 0 | 0 | Deceased | 10.1 |
| 160441 | 75 | M | No | No | II | 3 | 0 | 0 | Deceased | 19.6 |
| 160443 | 61 | F | No | No | III | 3 | 3 | 0 | Deceased | 19.3 |
| 160453 | 72 | M | Yes | Yes | II | 2 | 0 | 0 | Alive | 31.6 |
| 160458 | 66 | M | Yes | Yes | II | 3 | 0 | 0 | Deceased | 22.4 |
| 160460 | 59 | M | Yes | Yes | II | 3 | 0 | 0 | Alive | 30.6 |
| 160461 | 63 | M | Yes | Yes | II | 2 | 1 | 0 | Deceased | 9.8 |
| 160466 | 66 | M | Yes | Yes | III | 3 | 1 | 0 | Alive | 30.5 |
| 160467 | 73 | M | No | Yes | I | 1 | 0 | 0 | Alive | 30.3 |
| 160469 | 58 | F | No | No | I | 1 | 0 | 0 | Alive | 30.6 |
| 160473 | 48 | M | Yes | Yes | II | 3 | 0 | 0 | Alive | 30.6 |
| 160474 | 51 | M | yes | Yes | III | 3 | 1 | 0 | Alive | 30.6 |
| *M, male; F, female. ^#^Individuals who smoked an average of <1 cigarette/d and for <1 year in their lifetime were defined as nonsmokers; otherwise, they were defined as smokers. Individuals were classified as drinkers if they drank at least twice a week and continuously for at least 1 year during their lifetime; otherwise, they were defined as nondrinkers. ^§^Tumor TNM staging components, including tumor (T), lymph node (N) and metastasis (M), were reviewed by at least 3 pathologists and defined according to the American Joint Committee on Cancer (AJCC) 7th edition. | | | | | | | | | | |
