## Supplementary Table S3 for "Metabolic remodeling by TIGAR overexpression is a therapeutic target in esophageal squamous-cell carcinoma"

**Supplementary Table S3.** Small Interfering RNA Sequences for Expression Knockdown of the Investigated Genes in This Study

| Gene symbol | Small interfering RNA sequence (5' → 3') |
| --- | --- |
| *CCND1* | #1: GUUCGUGGCCUCUAAGAUG |
|  | #2: CCGAGAAGCUGUGCAUCUA |
|  | #3: GAACAGAAGUGCGAGGAGG |
| *CTTN* | #1: GAACAAGACCGAAUGGAUA |
|  | #2: GAAUAUCAGUCGAAACUUU |
|  | #3: GGACAGAGUUGAUCAGUCU |
| *PPFIA1* | #1: GAAGAAAGGUUACGACAGA |
|  | #2: GAACGACACUCGCCUAUGG |
|  | #3: GUGAUUCGCUGGAUCCUGU |
| *FADD* | #1: CAGCAUUUAACGUCAUAUG |
|  | #2: UGCAGCAUUUAACGUCAUA |
|  | #3: GUGCAGCAUUUAACGUCAU |
| *ANO1* | #1: GCAGAGAGGCCGAGUUUCU |
|  | #2: GCACGAUUGUCUAUGAGAU |
|  | #3: UUACGUGGCGUUCUUCAAA |
| *ORAOV1* | #1: GAAAUUCCCUUAUGAUGAC |
|  | #2: GCAGAAGGUUCCGGACUUU |
|  | #3: GAAGAAGGCAGUAGUUUGG |
| *ACTL6A* | #1: GGAAAUAGAUGGCGAUAAA |
|  | #2: GAACGGAGGUUUAGCUCAU |
|  | #3: GAUGCCAACUGUUCAUUAU |
| *FXR1* | #1: CCAUACAGCUUACUUGAUA |
|  | #2: GUAAACAUCUUAAGUGACA |
|  | #3: CGUACGAAGUUGAUGCUUA |
| *KCNMB3* | #1: UGAAGAGGCUGUCCAGAUA |
|  | #2: UCAGAAAGCUCUCCUACAU |
|  | #3: GAGACAGACUACAGUGAUG |
| *ZNF639* | #1: GAGCAUGCAUGUAAAUUAA |
|  | #2: CAACAAAGAUGAUGAUUCU |
|  | #3: GGAGUAGACUUCACACAAA |
| *MRPL47* | #1: GAAACGAGCCCGCAUCAAA |
|  | #2: GUCCAGAGCGGUUAGAUAA |
|  | #3: GUUAUACCUUGGCACCUAA |
| *NDUFS6* | #1: GUUUGUAGGUCGUCAGAAA |
|  | #2: GGUGGAGACUCGGGUGAUA |
|  | #3: ACACUGGCCAGGUUUAUGA |
| *MRPL36* | #1: GGUGGUACGUCUACUGUAA |
|  | #2: ACAAGACUGUCCUUAAGAA |
|  | #3: GGUGAACCCUCUGCUCUAU |
| *BRD9* | #1: CAAUGAAGAUACAGCUGUU |
|  | #2: GGCCAGAUACCGUGUACUA |
|  | #3: CACGCAGGCUUUAAGAUGA |
| *LPCAT1* | #1: GAACUCUGAUCCAGUAUAU |
|  | #2: GGACAGAUACUCAGAAAGA |
|  | #3: GGAAAGUGGCCACAGAUAA |
| *CLPTM1L* | #1: GCACUUACAGCGAAUCUGA |
|  | #2: GAAGAAAUCAACCUGCUCA |
|  | #3: GAAACAAUGGGACGCUGUA |
| *MCM7* | #1: GGAAAUAUCCCUCGUAGUA |
|  | #2: GGAGAGAACACAAGGAUUG |
|  | #3: UGUCAUACAUUGAUCGACU |
| *PMS2P1* | #1: GGACUUAACUGGAGAGGUU |
|  | #2: GAAAGGAGCUUCUGAUUGC |
|  | #3: CAAUGUAGCUCUGGAUAUA |
| *POLR2J* | #1: UGCGGACACUGGAAAGAAA |
|  | #2: CAGGAGACCAGGUGUCUGA |
|  | #3: GAAGAAAACCCGAGGUGAU |
| *TAF6* | #1: CUACCGAGCUUGCAAGUUU |
|  | #2: UGGCUGGAAUUUACUAUAU |
|  | #3: GGUAAGACUUUGUCAAGGG |
| *BUD31* | #1: GAAAGUGGAAUCUCUGUGG |
|  | #2: CUACAUCUUCGACCUCUUU |
|  | #3: GCGGAAAGCCAUCAGCAGA |
| *ATP5J2* | #1: CUUCAGUCCUAGUGGCAUU |
|  | #2: CAGUGAAGGACAAGAAACU |
|  | #3: AAACAUGACUGGUACGUGU |
| *POP7* | #1: CCAGUGGAAUACACCCUUA |
|  | #2: GAUCUACAUUCACGGCUUG |
|  | #3: ACAUGAAGACGGACUUUAA |
| *ARPC1A* | #1: ACGAAGUGCACAUCUAUAA |
|  | #2: GAAUUAAUCGCGCAGCUAC |
|  | #3: GUGGCACGAUGGCGAGGAA |
| *CPSF4* | #1: UGAAUUACCUCGUGGGAUU |
|  | #2: UUACAAGUGUGGCGAGAAA |
|  | #3: GCGCUGCUGUCUGUGAAUU |
| *PDAP1* | #1: CAAAAUAAAUUCCCGUGAA |
|  | #2: GCAAAAGCGCAAAGGCGUU |
|  | #3: CUGAGGAGAUCGACGCGCA |
| *BRI3* | #1: UGGAGCCACCUUCGCUUAA |
|  | #2: GCCUUGAGGAAGCGACGAU |
|  | #3: GCUCAUGACAACUCAAUAA |
| *PTCD1* | #1: AACCAGAGGUGGAUACUAA |
|  | #2: GAACUACACCUAUCUCAUC |
|  | #3: GGAUGUGCCUCGAUGUGUU |
| *PPP1R35* | #1: GCACAAAGAAAGUGAUUUA |
|  | #2: AAACUUCAGUCUCCGAUUA |
|  | #3: GAAAUUGACAGACGAAUAU |
| *LRWD1* | #1: GCAUCGUGCUCCACAAGUA |
|  | #2: GAAGUCGGCUGUCAGGGAU |
|  | #3: GAUUGGAGCUGCUUUCCGA |
| *C7orf59* | #1: GAACACACACUGCUGGUGA |
|  | #2: GCGCCUGUCUGUGGUCUUU |
|  | #3: GUGCAGUGCUGGCGUCAUC |
| *UFSP1* | #1: AGAGGUGAGUGCAGCCUUU |
|  | #2: GGUAUUGGACCCUCACUAC |
|  | #3: CUGCCUCGCUCACUUCGGA |
| *SPDYE3* | #1: GAGAAAGGAUGGCAAAGAU |
|  | #2: CAAACGGGCUGCAGAGAUG |
|  | #3: GAAACGAGCUCUGUGGAGU |
| *SEC61G* | #1: GCACUAAACCUGAUAGAAA |
|  | #2: GUAAAGGACUCCAUUCGGC |
|  | #3: GAUUUGCUAUAAUGGGAUU |
| *VOPP1* | #1: GAUGAACCCUGUCGGGAAU |
|  | #2: GAGCCAGCCUUCAAUGUGU |
|  | #3: GACUCUAUCCAACCUAUUA |
| *PSMA6* | #1: GAGCAAGGCCCUCAGGUAU |
|  | #2: CCUGACAAAUUAUUGGAUU |
|  | #3: CAAGUAUGGCUAUGAGAUU |
| *SRP54* | #1: GAAGAGGUAUUGAAUGCUA |
|  | #2: GAAGACCUGUUUAAUAUGU |
|  | #3: GAAAUGAACAGGAGUCAAU |
| *KIAA0391* | #1: GGGAAUCACUGCAGGUUUA |
|  | #2: AAACUAACCUUUCAGCGUA |
|  | #3: GGGAUAAACUUAAGGAAGA |
| *BAZ1A* | #1: CAAGUUAGAUUGCCAGUUA |
|  | #2: GGAAUUAGAUCAAGAUAUG |
|  | #3: GGUAUGAGCUGUUAAGUGA |
| *SNX6* | #1: GAUGAAGACCUCAAACUUU |
|  | #2: UAAAUCAGCAGAUGGAGUA |
|  | #3: CAAGAAGAGUUGCUGCAUU |
| *BRMS1L* | #1: UUAAUAGGCUCGUGGCUGA |
|  | #2: GGCACGAGCUGGCGUUUGU |
|  | #3: UGGCUGAGGUGGUCCGUAU |
| *SPTSSA* | #1: GCACUACUUUGAAAUCGUA |
|  | #2: GAGCGGACGGUGUUCAAUU |
|  | #3: AAGCAGAUGUCCUGGUUCU |
| *TIGAR* | #1: GGAAGGAAGAGAAGUUAAA |
|  | #2: CCUACAGGAUCAUCUAAAU |
|  | #3: GCUGGUAUAUUUCUGAAUA |
| *METTL4* | #1: GCAAUGAAGUUCUCAAAUU |
|  | #2: CUACGAAGGUCUUAUACUG |
|  | #3: GUUGUCAGCUGGGUGGUUA |
| *RTEL1* | #1: GACCAUCAGUGCUUACUAU |
|  | #2: GACAUUAUCCAGAUUGUGU |
|  | #3: UAUUCAUGCCGUACAAUUA |
| *YTHDF1* | #1: GAACAUGCCAGUUUCAAAG |
|  | #2: GGACAGUCAAAUCAGAGUA |
|  | #3: CGACAUCCACCGCUCCAUU |
| *ARFGAP1* | #1: ACAUUGAGCUUGAGAAGAU |
|  | #2: GGGAUAAGGUGGUCGCUCU |
|  | #3: GGGAGUCGGUAGUAAGGGA |
| *ZGPAT* | #1: GAAGAUGACUGAGUUCUAG |
|  | #2: CUACUACACAGUCAAGUUU |
|  | #3: UCAGAGACCGUUCCUAAAG |
| *ABHD16B* | #1: GCAACGAGAUCGACACUAU |
|  | #2: GAGAUGGGCUGUCUGUCUG |
|  | #3: GCACUUCAACCUCAACGUG |
| *POLB* | #1: GAAUUGGGCUGAAAUAUUU |
|  | #2: CAAGGAAGUUUGUAGAUGA |
|  | #3: GAUACGAGUUCAUCCAUCA |
| *AP3M2* | #1: GGACAGUAGUUCCCUUGGA |
|  | #2: CAAUGUAGUUGUGGUUUAU |
|  | #3: GUUCAGAGCCAGUGAUCAA |
| *CRKL* | #1: CCGAAGACCUGCCCUUUAA |
|  | #2: GAAGAUAACCUGGAAUAUG |
|  | #3: AAUAGGAAUUCCAACAGUU |
| *SMARCB1* | #1: GAAACUACCUCCGUAUGUU |
|  | #2: CCACAACCAUCAACAGGAA |
|  | #3: GUGACGAUCUGGAUUUGAA |
| *SNRPD3* | #1: GAAGAACGCACCCAUGUUA |
|  | #2: GAACACCGGUGAGGUAUAU |
|  | #3: CGAUUAAAGUACUGCAUGA |
| *UFD1L* | #1: AAUCAAGCCUGGAGAUAUU |
|  | #2: GACCAAACCCGACAAGGCA |
|  | #3: GAGCGUCAACCUUCAAGUG |
| *ZNF74* | #1: GAGCAAACCUCGAAACUAA |
|  | #2: GAACCACUGUCUCAUUAAA |
|  | #3: UGAAGGCGGUGCCCUCUCA |
| *TRMT2A* | #1: GAGAGAAGGCGCACCAUGA |
|  | #2: AGAGAUCAGUACAGUGUGA |
|  | #3: GAUAUUCCAGCCAGGGUAA |
| *MED15* | #1: CCAAGACCCGGGACGAAUA |
|  | #2: CGACAAGAACGAAGACAGA |
|  | #3: CGUCAGUGAUCCUAUGAAU |
| *MGC16703* | #1: GGACAAUGCUUUAAGUUUU |
|  | #2: CAAUAACUCUACUCAUUUG |
|  | #3: UCACAUCUAUAUCCAAGUU |
| *MAD1L1* | #1: GCUCUGGACUGGAUAUUUC |
|  | #2: AGAGGGAGCUUGCCUUGAA |
|  | #3: GAGCAGAUCCGUUCGAAGU |
| *INTS1* | #1: GGACAUCUAUUGCCUUUAA |
|  | #2: GACCACAGAUCGAUACGUA |
|  | #3: GGAAAUUACCGUUUACAAA |
| *YEATS4* | #1: GACAACAUCUCGUCAGCUA |
|  | #2: CAACAGCAAUGAUGCAACA |
|  | #3: UUACUAAACCUCCAUAUGA |
| *CCT2* | #1: GAAAUUGCCUCUACCUUUG |
|  | #2: AAAGUUAGCUGUAGAAGCA |
|  | #3: GGAGGAAGUUUGGCAGAUU |
| *CPSF6* | #1: GAAGAAAGCUUUGACAUCA |
|  | #2: GAAGAAGAAUGUACUACUA |
|  | #3: GGAGAUAUGUCUUUGUUAC |
| *NUP107* | #1: GGAAAUCUCUCCAUGGUUA |
|  | #2: GAAAGUGUAUUCGCAGUUA |
|  | #3: CAUCAGAGCUUAUUUGGAA |
| *RAB3IP* | #1: GGAUAAAGGGGUUGGGUAA |
|  | #2: GAACAUCGGUGGUGUAUUA |
|  | #3: GGAAAAUGUAAGACGUAAA |
| *PARP1* | #1: GAAAGUGUGUUCAACUAAU |
|  | #2: GCAACAAACUGGAACAGAU |
|  | #3: GAAGUCAUCGAUAUCUUUA |
| *ADSS* | #1: GAAACGAACCAGACCAAUA |
|  | #2: AAACGGAGGUCGUAUAAUG |
|  | #3: GGACAUCUCUAACAAACGG |
| *NVL* | #1: GAAACGAACCAGACCAAUA |
|  | #2: AAACGGAGGUCGUAUAAUG |
|  | #3: GGACAUCUCUAACAAACGG |
| *TARBP1* | #1: UCAGGGAUGUUAUUCAUUG |
|  | #2: UAAGAGAGAAGACCAUUAU |
|  | #3: CAGAGAAUGUAUUGCGGAU |
| *ZNF124* | #1: AAACUCAUAUUGCACAGAA |
|  | #2: CCAGUUCCCUUCAGAAACA |
|  | #3: CCAAUUGCCUUCAUUACCA |
| *MAPKAPK2* | #1: UCAAGGAACCAGAGAAUUA |
|  | #2: GUAUUACGGCUAUGUCAAA |
|  | #3: GUGGUAAUGAUCCAAAGUA |
| *CEP170* | #1: GAAGUAAAGUAACGAAAUC |
|  | #2: CGUAACAUCUCUCGGAUUG |
|  | #3: GAUUAUAAUAGGCCUGUUA |
| *COG2* | #1: GGACGGGCCUCGGGAGCUA |
|  | #2: UGGAGGAAGUGGCUUGCAA |
|  | #3: GGAGCCUGGUGUUGGUGAA |
| *ZNF281* | #1: GCACCACCGUGACGUAUUA |
|  | #2: CCAAUGACCUUUAUCACUA |
|  | #3: GCAGAAAGAAUACAGAUAA |
| *AHCTF1* | #1: GGGAGAAACUGCAGUACUA |
|  | #2: GGAUAGCGACCUGGAAAUA |
|  | #3: GGCGAAGGCAUCUCAGUUA |
| *FLVCR1* | #1: CGAAGACACAACAUAAAUA |
|  | #2: CCUGAAGAGUACUCCUAUA |
|  | #3: GGAUUGGGCUAACGCUAGU |
| *PYCR2* | #1: CCUGUCGGCUCACAAGAUA |
|  | #2: GCAACAAGGAGACGGUGAA |
|  | #3: GUCCAUGGCCGACCAAGAA |
| *PPPDE1* | #1: GCGCCACACUAAACUAUAA |
|  | #2: GAAAGAGAUUCCUCGCUGG |
|  | #3: AGAGUUGCCUCCCGAAGGA |
| *IPO9* | #1: UAUAAGAUCUUCACCAUGG |
|  | #2: GACAAACGGCUACAGGAUA |
|  | #3: CCUAAUGGGUUGAGAGAAU |
| *SMYD3* | #1: GGACAAACACGGACAGUAU |
|  | #2: GUACUAAGGUCUACAUCGA |
|  | #3: GGAGAGAAGCAGAAUAUUC |
| *TTC13* | #1: GCUAUGCUAUACAAAGGUU |
|  | #2: AAUGAAGUGUGCCAGUAUA |
|  | #3: CACUAAAGCUAUCCAACUG |
| *ZNF669* | #1: GUAAAGAAGAUGGUCAGUA |
|  | #2: GAUACAAACCAUAUGGAGA |
|  | #3: AAUCUUGAUCUGAACGAGA |
| *NTPCR* | #1: GCAUUGGGCUCAUGUUCUA |
|  | #2: GCAGUACCCUGAAGGACAA |
|  | #3: GACAUCCGCUUGACUCCUA |
| *ZNF670* | #1: UAAAGAAGGCAGUCAAUAU |
|  | #2: GAAAGAGCUUAUAUGUGGU |
|  | #3: UGUUGGAUCCCUAGACUAA |
| *SNAP47* | #1: GAAAGAAGGGAUACUGAUA |
|  | #2: GUUAGAAGAUGCAUUGGUG |
|  | #3: UAACAUCGCUGUCGCUCAG |
| *C1orf96* | #1: CCAAGUGCCUUAUUUGCUA |
|  | #2: GCGAGUACAUGAAGCGCUA |
|  | #3: AAACAGACCGAUACAGGAA |
| *MRPL55* | #1: GACAGAGUAUGCUCGCUAU |
|  | #2: UCAAGCACCUCCCUGAGAA |
|  | #3: GGACACUACUCUUACCGCA |
| *SYT14* | #1: UCACAGACAUCCCAACAUA |
|  | #2: GCAAACAGACCACCCAAUA |
|  | #3: CGGCAGAGUAUGAUGGAUA |
| *JAG2* | #1: GAACGGCGCUCGCUGCUAU |
|  | #2: GCAAGGAAGCUGUGUGUAA |
|  | #3: CCGGCCACCUGGACAAUAA |
| *TRAF3* | #1: GAUAAGGUGUUUAAGGAUA |
|  | #2: GACAUCUGCUGGUGCAUUU |
|  | #3: GGCCCAAACUGUUCUAGAA |
| *XRCC3* | #1: GGACCUGAAUCCCAGAAUU |
|  | #2: GCUGAGAACGGCCUCCUUA |
|  | #3: GGACCAGACUUGAAGAGAC |
| *CCNK* | #1: CAAAGCAACUCAAAGGUGA |
|  | #2: CAAGUUUGAUUUACAGGUA |
|  | #3: GAUCAUAGCAGUAGCAGUG |
| *MTA1* | #1: CCAUAGUGUUUGUCAUCUA |
|  | #2: GCCUGGAGGCGCACAAAUA |
|  | #3: GCGCACAAAUACCGGAACA |
| *SNHG10* | #1: GAGCAUGACUUCGUUAUUA |
|  | #2: GGAUCGGCCAGCUGAAUUA |
|  | #3: GGGAGUCGGUGAAGUACUU |
| *TOR1A* | #1: CAUCUUAAAUGCCGUGUUU |
|  | #2: AGAAAGCCAUGUUCAUAUU |
|  | #3: CUUCAAACAUCACCUUGUA |
| *FPGS* | #1: GACCAAGGAUGGCAGCUGU |
|  | #2: GAAUGUAUCCUCCGAAGCU |
|  | #3: GAACAUCAUCCACGUCACU |
| *GLE1* | #1: GGAAGGAGCUAAGUGCAAU |
|  | #2: GGAACUACUUCUCCCUGUA |
|  | #3: GCUCGGAUGUGCAGAACAA |
| *ODF2* | #1: GAAGGGAGACCGAGACAAA |
|  | #2: CAAAUGACCUGCACGGACA |
|  | #3: GCGGUUAAGUGACCUCUCU |
| *PSMB7* | #1: GGAAGAAACAGUCCAAACA |
|  | #2: CAUCGCAGCUGGCAUCUUC |
|  | #3: CCGCAGGAAUGCCGUCUUG |
| *SET* | #1: GAGGAAGGAUUAGAAGAUA |
|  | #2: GGAUGAAGGUGAAGAAGAU |
|  | #3: GAAGUCCACCGAAAUCAAA |
| *SNAPC4* | #1: GGACUUGGAUCCUGCCGAU |
|  | #2: AAGGGUCGGUGGAAUUUAA |
|  | #3: AAACGUGCCGGCUCAAGAA |
| *SURF4* | #1: GCAGGAACUUCGUGCAGUA |
|  | #2: GGACUUCGCCGACCAGUUC |
|  | #3: GCUCUUUGCCAUCAACGUA |
| *VAV2* | #1: UCACAGAGGCCAAGAAAUU |
|  | #2: GAAAGUCUGCCACGAUAAA |
|  | #3: GGGACGACAUCUACGAGGA |
| *GTF3C5* | #1: GCAGAUGUUCUACCAGUUA |
|  | #2: GCAAGCAUACGUCAAUGUA |
|  | #3: CGAAUCCGUUGUGGAAUGA |
| *MED27* | #1: GUAAAGGGAUAUAACGAGA |
|  | #2: UGAAAGUGAUCGUCGUCAU |
|  | #3: UGAAGGAUGGGAUGCGGAA |
| *RABEPK* | #1: CCACAGCUGUUCAUAUUUA |
|  | #2: GGAUUCAGCUGACAAAGUA |
|  | #3: GAAACCAGCUAUAUGUCUU |
| *SDCCAG3* | #1: GAACCUCGGCCUCUCGAAA |
|  | #2: GAGCGGAAGUUAGAAGCAA |
|  | #3: CCUCGUGGGCGUUGAGUGA |
| *WDR5* | #1: GAGAGUGGCUGGCAAGUUC |
|  | #2: GACGAAAGCGUGAGGAUAU |
|  | #3: CAGAGGAUAACCUUGUUUA |
| *MAN1B1* | #1: CUGCGAAGGUGGAGAGUUA |
|  | #2: GAAAGAGUCAUCAGAAGUG |
|  | #3: GGGAGUCGCUGCAAGAAUC |
| *DOLK* | #1: GAGAUCCGCUGGCCUGGAA |
|  | #2: CUACAGUUAUGCUUGGAUU |
|  | #3: CGCAAACAGUGGCCUAUUG |
| *EXOSC2* | #1: GAACGUAUAUGGGAGAAGA |
|  | #2: GAAACACGCCAGAGGCUUU |
|  | #3: GGGAUACAAUCACUACGGA |
| *COBRA1* | #1: GCCCUGAGGUGUUUCUGAA |
|  | #2: GCAGUUCAGUGUCAUCGUC |
|  | #3: UGACGCGACUGAUCUCAUG |
| *NELF* | #1: GAACGUCUACCACAAGGGA |
|  | #2: GUACAGCGUUGACCGUGUG |
|  | #3: CACCAAAGGUCAUGCUCAU |
| *TOR1B* | #1: GCAAUAAUGUAGACUGAUA |
|  | #2: UGACAACCCUCGAAGAUAU |
|  | #3: GGACUAAAGUAUUCCACAA |
| *METTL11A* | #1: GCAAGAGGGUGAGGAACUA |
|  | #2: GGAAGUUUCUGCAGAGGUU |
|  | #3: GCCAAGACCUACUGGAAAC |
| *MRPS2* | #1: CAUGAGCCAUUCCCUGUGA |
|  | #2: CGGCAGUUCUCGUACCUGA |
|  | #3: CCACAUGGCCUACCGCAAG |
| *C9orf114* | #1: GGAAGGAUCUCAAGCUGAU |
|  | #2: GCAGGACCCUCGCACCAAA |
|  | #3: CUACUUGGCCGGUCAGAUU |
| *REXO4* | #1: GGAAGUGCGUUUAUGACAA |
|  | #2: GCGGAGCACUGUUCAAUUC |
|  | #3: GAGGGGACAUCGAGCAUAA |
| *DDX31* | #1: UCGGUUAGCACAAGUGAUA |
|  | #2: GGAGAGACAGUGCAUUAAG |
|  | #3: CGCCGGAUCUCGCUUCUCA |
| *C9orf69* | #1: ACAACAACUUCGACGUGUA |
|  | #2: GGUCUUCGUGGACAUGUGA |
|  | #3: UCAAGACGCGCUGGCUGUA |
| *UAP1L1* | #1: UGAAGAAGGUCCCGUAUGU |
|  | #2: CCAACGUGGUCAUGUUUGA |
|  | #3: GAGAGGGUUUAGAAGUGUA |
| *WDR85* | #1: CAGGGUACCCGGCAAAUUU |
|  | #2: GCAGAGAGGAUAACGAUGG |
|  | #3: GAAGGGUGCAAGCGAGUUG |
| *ZMYND19* | #1: GAAUGGAAGUGGAUGCAGA |
|  | #2: CUACAGACCCUAUAGAAGA |
|  | #3: CUAAAUGUGACCCGGUAUU |
| *PTRH1* | #1: GGAGUCCGUUCCUGCAUUA |
|  | #2: GCCAUGAGCCGAUGUGUUU |
|  | #3: GACCUGAUCUUGGACCACA |
| *QSOX2* | #1: GACCUGAUCCCGUAUGAAA |
|  | #2: GAUGCGGAUUUCUGGAAUA |
|  | #3: UAAAGUGGGUUGGAUGUCA |
| *C9orf142* | #1: GGACAGAGCAUCCCUGACG |
|  | #2: CAGGAGAGUCGCUCAUCAA |
|  | #3: GAGUGCGGCUGAGGACAUC |
| *DNLZ* | #1: CGGAAGCGGGUGAGGAUGA |
|  | #2: CUGCAAGGUCUGCGGGACU |
|  | #3: GAAGAGAAAUAUCGAAGAG |
| *LOC100272217* | #1: CAGACUACUUGCUACAUUA |
|  | #2: CAAGUACUAUCGCCAGACA |
|  | #3: AAACUACCCACGAGCUUAA |
| *SSB* | #1: GAAAUGAAAUCUCUAGAAG |
|  | #2: GAACAUUGCAUAAAGCAUU |
|  | #3: AGAUAAAGGUCAAGUACUA |
| *HAT1* | #1: GCACAAACACGAAUGAUUU |
|  | #2: GAAGAUUACCGGCGUGUUA |
|  | #3: GCUACAGACUGGAUAUUAA |
| *OLA1* | #1: CAACAAGGCAGAAAUUAUA |
|  | #2: UAACACACCUCAACAACCG |
|  | #3: AGAAGUAUCUGGAAGCGAA |
| *ABCF1* | #1: GAUGAUGUCUGCACUGAUA |
|  | #2: GGACAGAAACCACUCUUUA |
|  | #3: CCAAAUCGAUGGUGACUUU |
| *GTF2H4* | #1: GAAACGGAGUCUAAGCUUU |
|  | #2: GCUCUGAAGUGCAUCAAGA |
|  | #3: GAACAGAAUGGCAUCAUGU |
| *NEU1* | #1: GAAAGAGACAGUCCAGCUA |
|  | #2: CGGAAUCUCUCCCUGGAUA |
|  | #3: CGAUGGAGCUUCAGCAAUG |
| *SKIV2L* | #1: GCUACAAGAACGUGAAGAA |
|  | #2: GACCUCACCUGGUACGCCA |
|  | #3: GGGAGAACUACCUGGGCAA |
| *RDBP* | #1: GAAGAUAUCCGGUGCAAAU |
|  | #2: GAAUGAGGAUGCUCGCUCU |
|  | #3: GAAAGCCGGAUGCAUAUAC |
| *DHX16* | #1: GGCAGGAGCUCAAAUAUAA |
|  | #2: GAGAGUACCUGGCUAAGCG |
|  | #3: GAACUGUCCUCCGCUACAU |
| *MDC1* | #1: GGACCAAACUUAACCAAGA |
|  | #2: GAAGACAACUAUGGUGAUU |
|  | #3: GAGCCCACAUCUCAGGUUA |
| *EHMT2* | #1: GAAGAGAGAUCAGCAUUAA |
|  | #2: GAACAAGGACCUAAACAAA |
|  | #3: GCAUAAAGUUCAACAGUAU |
| *NRM* | #1: CUUCAGAGGUCACUGUAUG |
|  | #2: CCAAAGGCCCUGUGUUGUG |
|  | #3: GCAUGGACAUCCCGGUACU |
| *MRPS18B* | #1: CAAGGAACCUGGUGUUGAA |
|  | #2: GGAAUGGAAAGUAGGAUUA |
|  | #3: CAAAUCUUCUUCUGAACUA |
| *CCHCR1* | #1: GAGACAACCUUGACAGAUG |
|  | #2: UUAAGCAGCUGAAGGGACA |
|  | #3: CCUGAUUGCUCGAAAGCUU |
| *PRR3* | #1: GGAGAGAGACGACCGUGUA |
|  | #2: GAGAGAAGGCCACUGAUGA |
|  | #3: GUACCCACCUGAGCUAUUC |
| *ZNF146* | #1: GCAAAUCCAACCUUACUGA |
|  | #2: AGAAGUACCUCAUAAAACA |
|  | #3: GCUCAGGAAAUACGCACUA |
| *UBA2* | #1: GCCGUUAGCCUGAUCAUCU |
|  | #2: AGCAUGUCAUCCACAAGUA |
|  | #3: CCUAUGUGAUGCUGAAGAC |
| *HAUS5* | #1: GAGGAAAGCCAAAGUAGAU |
|  | #2: UCAAGGCCCUGCACGAUCA |
|  | #3: GAAACCUACUCUGGUAUGG |
| *ALKBH6* | #1: ACAAGAACCUCCUGGUGUC |
|  | #2: CGUCCAACAUCUCAAACUU |
|  | #3: GCACCAGCACCUAGCGUAC |
| *RHPN2* | #1: GAGGAGGCAUUUACGAUUC |
|  | #2: GCUUAGCCAUUGAUGAUGA |
|  | #3: CGGAGUAAAUUGCAGAAUC |
| *GLS* | #1: CCCUGAAGCAGUUCGAAAU |
|  | #2: CCAGGUUGAAAGAGUGUAU |
|  | #3: GAGGCAUUCUACUGGAGAU |
| *PRKAA1* | #1: GAGGAGAGCUAUUUGAUUA |
|  | #2: GCGUGUACGAAGGAAGAAU |
|  | #3: CGGGAUCAGUUAGCAACUA |
